# Site saturation mutagenesis of 500 human protein domains reveals the contribution of protein destabilization to genetic disease

**DOI:** 10.1101/2024.04.26.591310

**Authors:** Antoni Beltran, Xiang’er Jiang, Yue Shen, Ben Lehner

## Abstract

Missense variants that change the amino acid sequences of proteins cause one third of human genetic diseases^1^. Tens of millions of missense variants exist in the current human population, with the vast majority having unknown functional consequences. Here we present the first large-scale experimental analysis of human missense variants across many different proteins. Using DNA synthesis and cellular selection experiments we quantify the impact of >500,000 variants on the abundance of >500 human protein domains. This dataset, Human Domainome 1, reveals that >60% of pathogenic missense variants reduce protein stability. The contribution of stability to protein fitness varies across proteins and diseases, and is particularly important in recessive disorders. Combining stability measurements with protein language models annotates functional sites across proteins. Mutational effects on stability are largely conserved in homologous domains, allowing accurate stability prediction across entire protein families using energy models. Domainome 1 demonstrates the feasibility of assaying human protein variants at scale and provides a large consistent reference dataset for clinical variant interpretation and the training and benchmarking of computational methods.

## Introduction

The human genome encodes >20,000 proteins. Missense variants in nearly 5,000 of these proteins cause Mendelian diseases^2^, however the functional consequences of nearly all missense variants in nearly all proteins are unknown^3–5^. Given the current human population size, most variants compatible with life are present in someone currently alive^6–8^, making the large-scale experimental analysis of variant function a central challenge for human genetics^7–9^. However, experiments to date have mostly quantified variant effects in one or a few proteins^7,10^. Despite recent improvements^11–14^, computational variant effect predictors are not deemed to provide sufficient evidence to classify clinical variants as pathogenic or benign^15^. They also do not identify the molecular mechanisms by which variants cause disease, information that is important for therapy development and clinical trial design. Whereas many disease variants are likely to destabilize proteins and reduce their abundance^8,16,17^, others may affect specific molecular interactions, or cause gain-of-function phenotypes^18,19^.

Previous studies have established reduced abundance as a frequent causal mechanism for pathogenic variants in diverse proteins^8,20–24^, but larger scale studies of human disease variants across many disease genes to test the generality of these observations are lacking. Most human proteins contain multiple independently folding structural units called domains^25,26^. For example, the human genome encodes more than 200 homeodomains that bind DNA to control gene expression and more than 250 PDZ domains that mediate protein-protein interactions^27,28^. The small size of protein domains (median ∼100 amino acids) and their independent folding make them a useful target for large-scale experimental measurement of variant effects^29^.

Here, using a highly-validated assay that quantifies the effects of variants on protein abundance in cells^30^ we perform the first large-scale mutagenesis of human protein domains. In total we report the impact of >500,000 missense variants on the stability of >500 different human domains. This dataset, which we refer to as Human Domainome 1, provides a large reference dataset for the interpretation of clinical genetic variants, and for benchmarking and training computational methods for stability variant effect prediction. We use the dataset to quantify the contribution of stability changes to human genetic disease and how this varies across proteins and diseases. We also show how stability measurements can be combined with protein language models to annotate functional sites across proteins, and that measurements made on a small number of proteins can be used to accurately predict stability changes across entire protein families.

## Results

### Massively parallel saturation mutagenesis of human protein domains

We used mMPS synthesis technology (manuscript in preparation) to construct a library of 1,230,584 amino acid (aa) variants in 1,248 structurally diverse protein domains (Extended Data Table 1). In this library, which we refer to as Human Domainome 1, every amino acid is mutated to all other 19 aa at every position in each domain (Fig. 1a). Sequencing the library revealed it to be high quality, with coverage of 91% of designed aa substitutions (Extended Data Table 1, Extended Data Fig. 1a). To quantify the impact of these variants on domain stability, we used an in cell selection system, abundance protein fragment complementation assay (aPCA)^30,31^. In aPCA, the protein domain of interest is expressed fused to a fragment of an essential enzyme and its concentration linearly determines the cellular growth rate over at least three orders of magnitude^30^ (Fig. 1b). The effects of variants on protein abundance are quantified using high-throughput sequencing to measure the change in variant frequencies between the input and output cell populations in selection experiments (Fig. 1a). This strategy thus allows pooled cloning, transformation and selection of hundreds of thousands of variants in diverse proteins in a single experiment (Fig. 1a).

**Figure 1:**
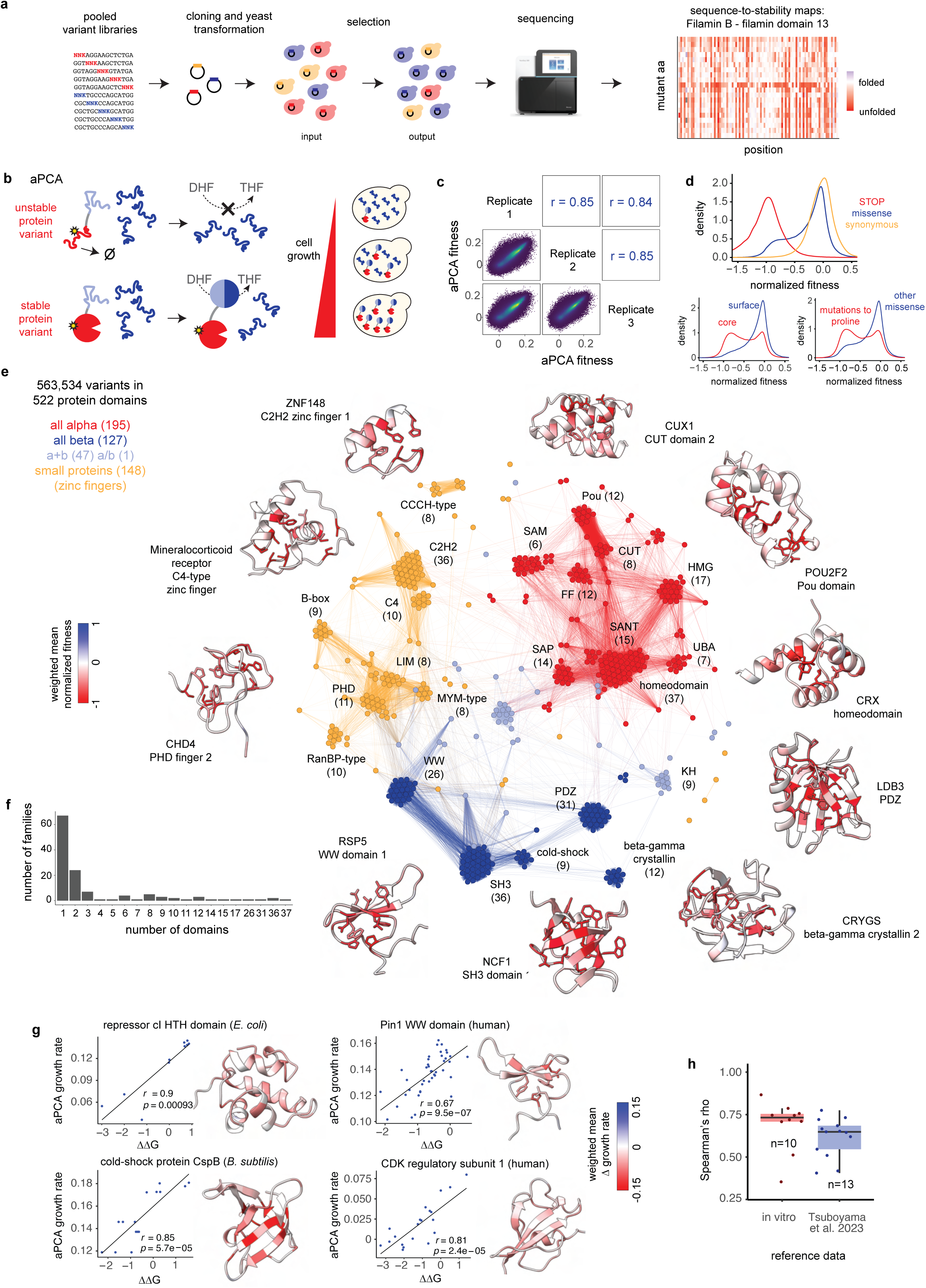
Mutating the Human Domainome. **a.** Experimental strategy for multiplexed generation of sequence-to-stability maps of human protein domains based on pooled cloning, transformation and selection of pooled saturation mutagenesis libraries. **b.** DHFR complementation assay to measure in vivo stability of variant human protein domains (abundancePCA^30^, aPCA). Variants causing unfolding and degradation of the target domain result in a decrease in concentration in the cell, leading to impaired cell growth. **c.** Replicate correlations of aPCA fitness scores. **d.** Comparison of aPCA fitness score distributions of synonymous, missense, and nonsense variants (top panel), core (rSASA < 25%) and surface variants (bottom left panel), and mutations to proline and other missense variants (bottom right panel). **e.** Structural similarity network of domains retained in the final dataset. Nodes represent protein domains, and edges represent foldseek hit probabilities in pairwise structural alignments. Colors correspond to SCOP structural classes. **f.** Distribution of number of domains per family in the library. **g.** Correlations between in vitro ΔΔGs and aPCA scores in four representative domains. **h.** Distribution of correlations (Spearman’s rho) between aPCA scores and in vitro ΔΔGs^32,33^ or stabilities derived by a high-throughput proteolysis assay^29^.

In total, we performed 27 transformation, selection and sequencing experiments (three independent replicates of nine sub-libraries). After filtering (see Methods), the final dataset consists of cellular abundance measurements for 563,534 variants in 522 protein domains, of which 503 are human (Extended Data Table 2). Abundance measurements were highly reproducible (median Pearson’s correlation coefficient, r = 0.85 between replicates for all variants, Fig. 1c, Extended Data Fig. 1b). They also correlated well to independent *in vitro* measurements of protein fold stability^32,33^ (median Spearman’s rho = 0.73 with folding free energy changes (ΔΔG), n=10 domains, Fig. 1g,h, Extended Data Fig. 1d). Moreover, they correlated well with high-throughput stability measurements from protease sensitivity assays^29^ (median rho = 0.65, n = 13 domains, Fig. 1h, Extended Data Fig. 1e).

The 522 domains are structurally diverse, constituting 195 all alpha, 127 all beta, 47 alpha+beta, 1 alpha/beta, and 148 metal-binding zinc finger domains (Fig. 1e). In total, they cover 127 different domain families, including 14 families with 10 or more homologous domains (Fig. 1f, Fig. 2), and 97 families with only one or two domains (Fig. 1f). Altogether they comprise 2.1% of all proteins, 1.2% of all domains and 2.0% of all unique domain families in the human proteome. 275 of the domains are encoded by human disease genes with 108 domains containing annotated pathogenic variants. Across the dataset, mutations in the buried cores of the domains are more detrimental than in their surfaces (Fig. 1d), with mutations to polar amino acids having stronger destabilizing effects in cores and mutations to hydrophobics having stronger effects in surfaces (Extended Data Fig. 1f). Mutations to proline are the most detrimental overall (Fig. 1d), both in core and surface residues, with highly destabilizing effects in beta strands and helices and milder effects in coils (Extended Data Fig. 1f,g). The full mutagenesis dataset is presented in Extended Data Fig. 2.

**Figure 2:**
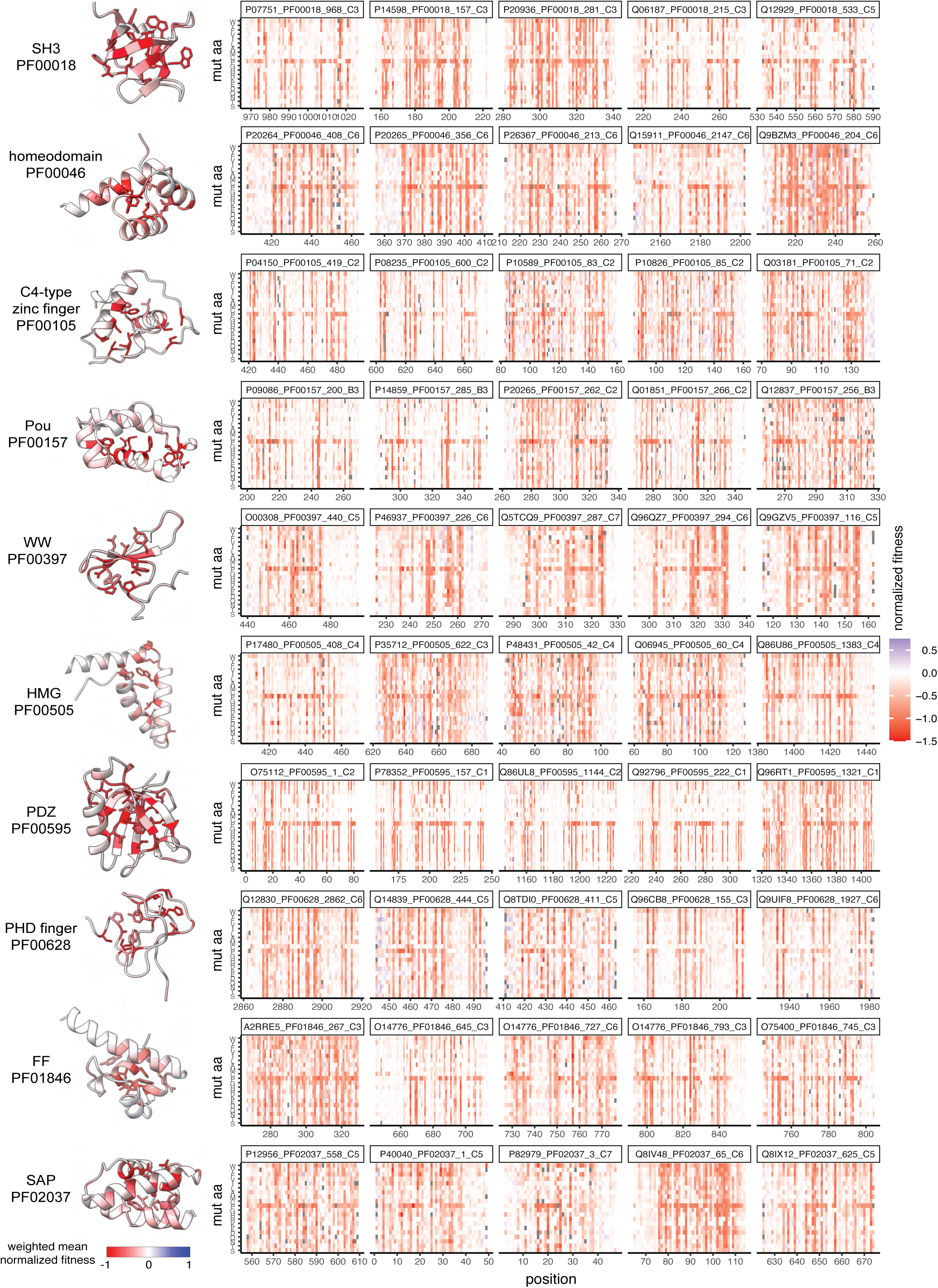
Deep mutational scans of protein homologs. Five examples of deep mutational scanning aPCA datasets of the most abundant protein families. Heatmaps depict the effects of mutating every residue in the protein domains (x axis) to all possible 19 amino acids (y axis).

### Evaluation of variant effect predictors on protein abundance

Our dataset represents a nearly 10-fold increase in stability measurements for mutations in human protein domains (Fig. 3a) and so an opportunity to evaluate how well computational variant effect predictors (VEPs) predict changes in stability. Several general VEPs provide reasonable prediction of abundance changes, including the protein language model ESM1v^12^ (median rho = 0.48) and the deep generative model EVE^11^ (median rho = 0.48). Amongst dedicated stability predictors, the graph neural network ThermoMPNN^34^ performs best (median rho = 0.50, Fig. 3b,c), and is the best predictor overall. Interestingly, all tested stability predictors perform poorly on small zinc finger domains that require metal binding for stability (Extended Data Fig. 3). After excluding zinc fingers, ThermoMPNN is still the best performing method overall (median rho = 0.57, Extended Data Fig. 3), even when evaluating on protein domains with no homology to the Megascale dataset used for ThermoMPNN training (median rho = 0.57, Extended Data Fig. 3).

**Figure 3:**
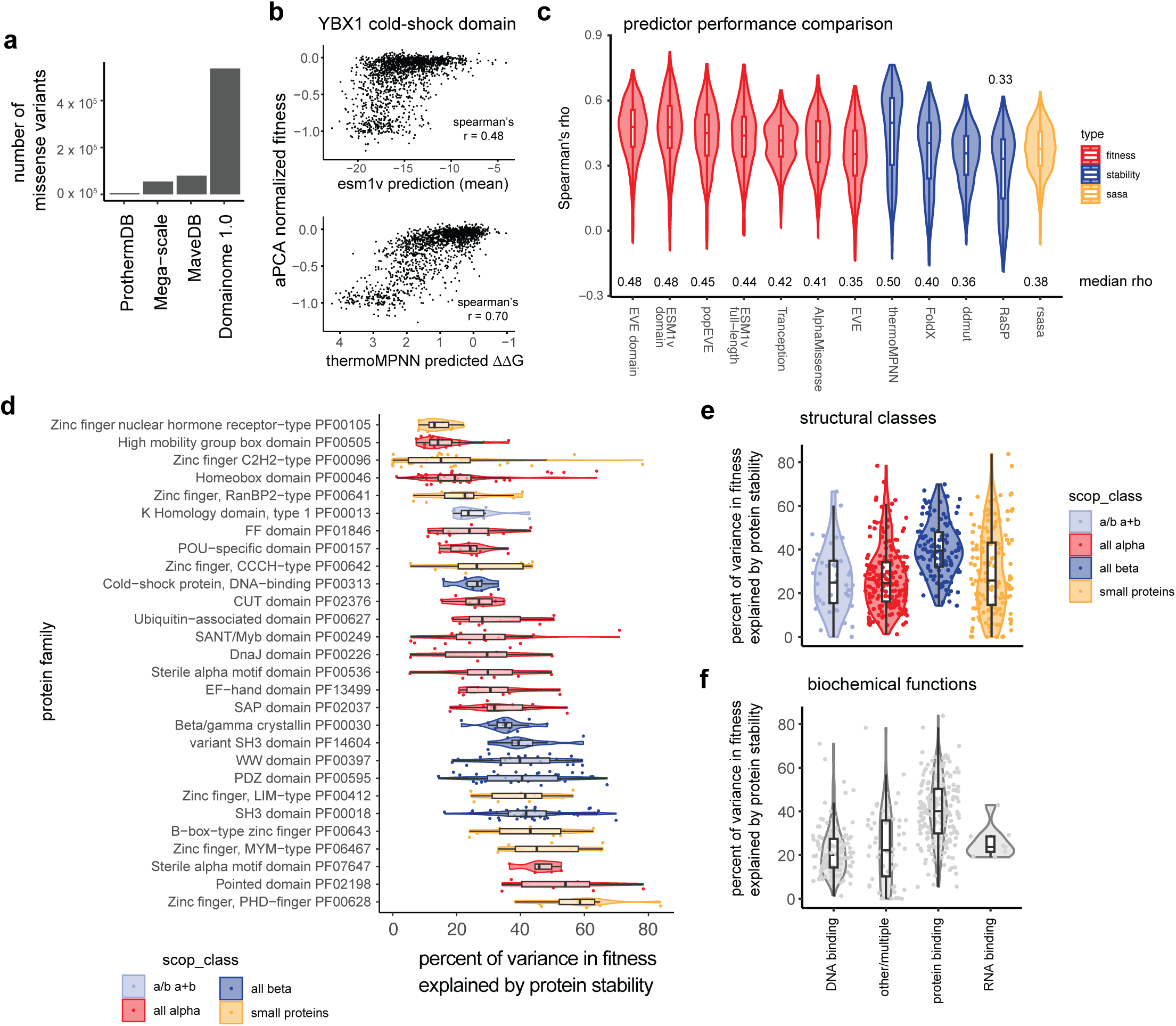
The contribution of protein stability to evolutionary fitness. **a.** Total number of stability measurements for human single missense variants in the Domainome 1 compared to previous comprehensive datasets of protein stability^29,32,62^. **b.** Correlations between aPCA normalized fitness and ESM1v predictions (top panel) or ThermoMPNN stability predictions (bottom panel) for the YBX1 cold-shock domain. **c.** Comparison of performance of variant effect and protein stability predictors. The distribution of Spearman’s r across all domains is shown for each predictor, with the median Spearman’s r labeled in text. **d.** Quantification of the degree of sequence constraint attributable to protein stability across protein families with at least 5 measured homologs. Colors correspond to SCOP structural classes. **e-f.** Quantification of the degree of sequence constraint attributable to protein stability across domains grouped by their SCOP structural classes (**e**) and molecular functions (**f**).

### The contribution of stability to protein fitness

Fold stability is just one of many biophysical properties that contribute to protein function and protein sequence conservation during evolution. In particular, many proteins bind other proteins, nucleic acids and small molecules or catalyze enzymatic reactions, and these functions often trade off with stability^35^. The extent to which selection on protein sequences is driven by changes in stability rather than other biophysical properties is an important open question^36^.

To address this, we compared experimentally quantified stability to evolutionary fitness quantified by ESM1v across >500,000 variants in >500 domains (Methods). We find that protein stability accounts for a median of 30% of the variance in protein fitness across all domains (Fig. 3d). However, the contribution of stability to fitness varies across domain families, with stability making a larger contribution to the fitness of all-beta domains (40% of the variance) than to that of all-alpha (25%) and mixed domains (25%) (Fig. 3e). This is consistent with the lower structural tolerance to mutations of beta sheets compared to alpha helices^37^, and suggests that stability is a more important determinant of the fitness landscapes of all-beta domains. The overall contribution of stability to protein fitness also varies in domains with different molecular functions. For example, stability has a larger contribution to fitness in SH3, WW, PHD finger, and Pointed protein-protein interaction domains, and a lower contribution in DNA-binding HMG-box domains, nuclear hormone-type C4 zinc fingers, and homeodomains (Fig. 3f).

### Identification of functional sites

Mutations in binding interfaces, active sites and allosteric control sites typically have larger effects on function than can be accounted for by changes in protein stability^23,25,31^. For example, quantifying the effects of mutations on protein binding and abundance allows binding interfaces and allosteric sites to be comprehensively mapped^25,37^. We reasoned that a similar approach could be used to identify functional sites in hundreds of domains by combining our abundance measurements with evolutionary fitness quantified by ESM1v.

For individual domains, protein abundance is non-linearly related to evolutionary fitness predicted by ESM1v (Fig. 4a, Extended Data Fig. 4a). We used sigmoidal curves to model this relationship (n = 426 domains, ESM1v fitness range > 10, WT aPCA fitness percentile < 30, Fig. 4a, Extended Data Fig. 4a). The residuals to these fits identify mutations with larger or smaller effects on evolutionary fitness than can be accounted for by changes in stability (Fig. 4a,b).

**Figure 4:**
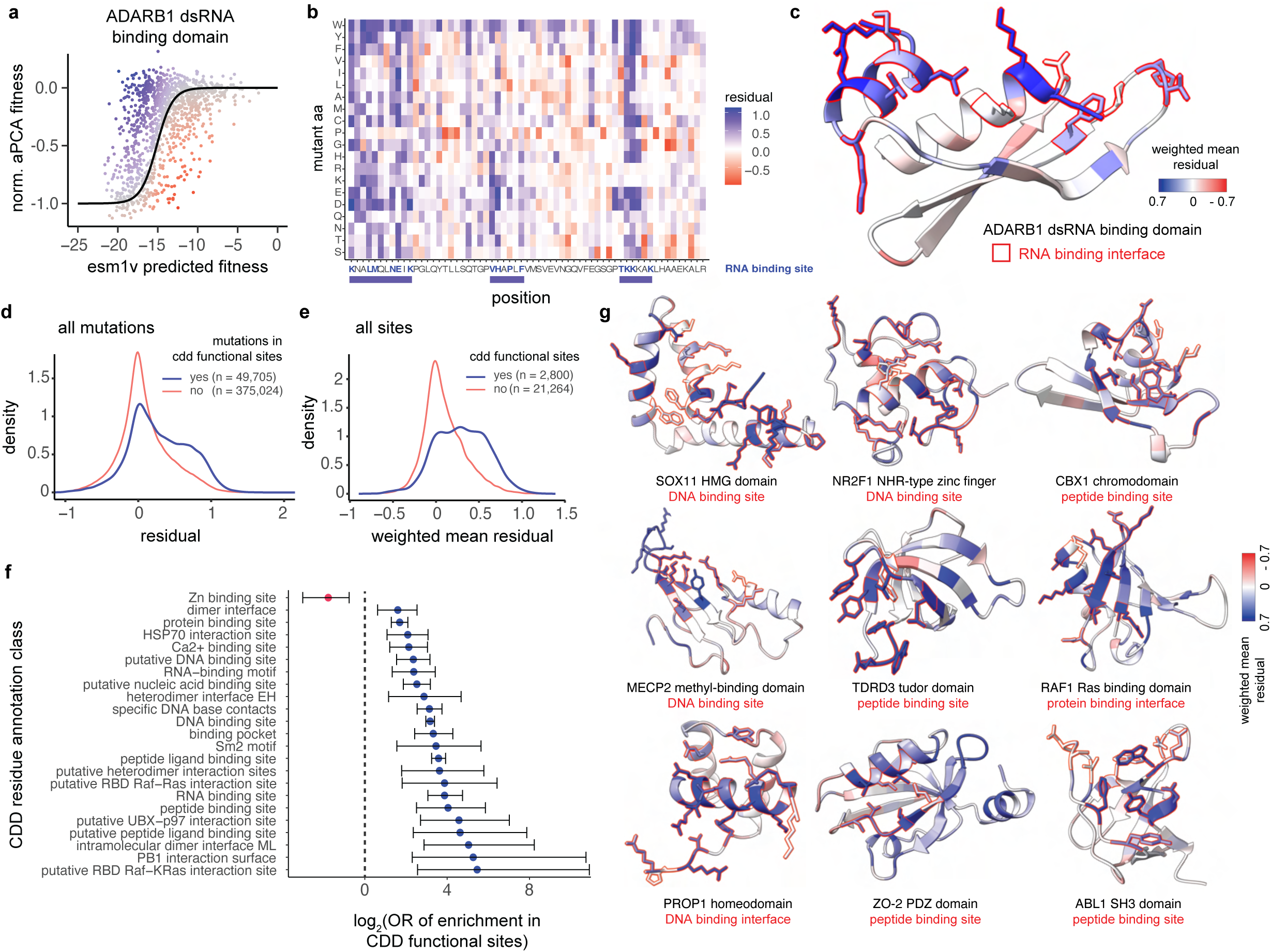
Identification of functional sites. **a.** Sigmoidal curves to model the relationship between stability and overall fitness of variants in the dsRNA binding domain of ADARB1. Data points are colored by their residuals to the fit. **b.** Heat map depicting the residuals to the fit of all measured variants. The dsRNA binding site residues are marked with blue bold letters. The blue boxes capture all the contiguous stretches of sequence containing the binding site. **c.** AlphaFold2 predicted structure of the dsRNA binding domain of ADARB1, with residues coloured by the weighted mean residuals to the fit. Residues forming the dsRNA binding site are marked with a red silhouette. **d.** Distribution of residuals for mutations in annotated functional sites, and in other protein residues. **e.** Distribution of weighted mean residuals of protein residues in annotated functional sites, and of other residues. **f.** Enrichment of functional sites (residues with weighted mean residuals > 0.3) in several types of CDD annotated functional sites. Error bars depict the 95% confidence interval for the odds ratio of enrichment. **g.** AlphaFold2 predicted structures of representative domains containing functional sites, with residues coloured by the weighted mean residuals. Residues that belong to CDD functional sites are marked with a red silhouette.

This analysis identifies a total of 102,231 mutations with larger effects on evolutionary fitness than can be accounted for by changes in stability (24% of the total; Z-test FDR < 0.1, normalized aPCA residual > 0.3). These mutations are enriched in known functional sites annotated in the conserved domains database (CDD, OR = 2.72, FET p < 2.2e-16, n = 3,104 functional sites in 2,800 residues, Fig. 4d, Extended Data Fig. 4b). Defining evolutionary functional sites as residues with a weighted mean residual > 0.3 identifies a total of 5,231 sites in 426 domains. These sites are strongly enriched in CDD annotated sites (OR = 4.50, FET p < 2.2e-16, Fig. 4e) and identify many known DNA, RNA, and protein binding interfaces (Fig. 4e,f). However, these evolutionary functional sites also include 1,942 sites in 180 domains without existing CDD annotations, and 1,873 additional sites in domains with other CDD annotations (Extended Data Table 3).

Interestingly, evolutionary functional sites without known annotations are located in closer proximity to annotated functional sites (median side chain heavy atom distance = 3.62 Å) than the rest of residues (median d = 6.93 Å, p < 2.2 x 10^-16^). Many of these therefore act as ‘second shell’ residues contacting DNA, RNA and protein binding interfaces (Fig. 4c,g, Extended Data Fig. 4c), where sequence changes may indirectly impact binding via energetic interactions with interface residues^25,37^.

### The contribution of stability to pathogenicity

Domainome 1 contains 3,652 variants with clinical annotations. 621 of these are classified as pathogenic/likely pathogenic (henceforth pathogenic), 322 as benign/likely benign (henceforth benign) and 2,709 as variants of uncertain significance (VUS), with 116 domains containing at least one pathogenic variant. Pathogenic variants are unevenly distributed across domains, with 75% of pathogenic variants contained in 25% of domains, and 41 domains containing only a single pathogenic variant (Extended Data Fig. 5a).

380/621 pathogenic variants (61%) cause a detectable domain destabilization (Z-test, FDR < 0.1) and 303/621 (48%) are strongly destabilizing (FDR < 0.1, normalized aPCA fitness < -0.3). This contrasts with 129/322 (40%) and 50/322 (16%) of benign variants, respectively. However, the association between pathogenicity and destabilization varies across domain families (Fig. 5b, Extended Data Fig. 5b). For example, many pathogenic mutations in beta-gamma crystallins that cause cataract disease are strongly destabilizing (13/18, odds ratio (OR) = 11.98 compared to benign variants, p = 1.99 x 10^-6^, Fisher’s exact test (FET), Fig. 5b). In contrast, a smaller proportion of pathogenic variants are strongly destabilizing in homeodomains (OR = 3.94, p = 1.45 x 10^-7^), HMG-box domains (OR = 2.10, p = 0.045), and CUT domains (OR = 1.65, p = 0.55) (Fig. 5b), all of which bind DNA, suggesting that many pathogenic variants in these domain families affect other biophysical properties such as DNA binding. Consistent with this variable association of destabilization with pathogenicity, aPCA has an overall lower performance than variant effect predictors in the classification of clinical variants across all domains, as the readout is specific for protein stability (Extended Data Fig. 5c,d).

**Figure 5:**
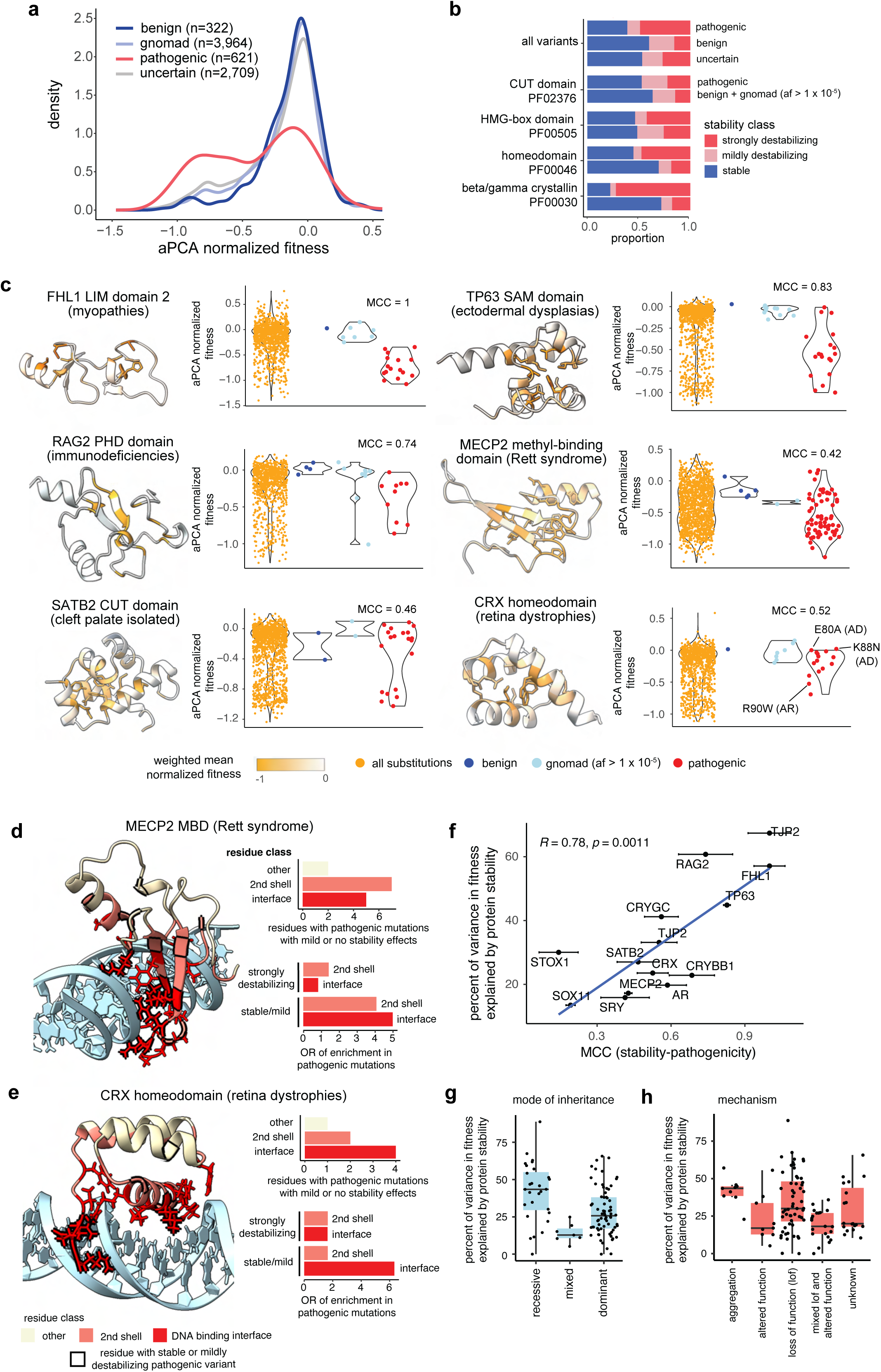
The contribution of destabilization to genetic disease. **a.** Distributions of normalized aPCA fitness values of pathogenic, benign, uncertain, and gnomAD variants (allele frequency > 10^-5^). **b.** Proportions of stability classes (stable, mild, strongly destabilizing) in the full dataset (top bars), and in several protein families. **c.** Distributions of mutational effects on stability in representative examples of human disease protein domains, showing all measured variants (yellow), benign variants (dark blue), gnomAD variants with an allele frequency > 1 x 10^-5^ (light blue), and pathogenic variants (red). Specific CRX variants with their associated mode of inheritance are labeled. AD = autosomal dominant, AR = autosomal recessive. MCC = Matthew’s correlation coefficient. **d-e.** Spatial distribution of mutations that do not strongly destabilize the MBD domain of MECP2 (d) and the CRX homeodomain (e). **f.** Correlation between the classification performance of stability on clinical variants (measured using Matthew’s correlation coefficient, MCC), and the fraction of protein fitness explained by stability changes, for domains with at least 20 clinical and gnomAD variants (allele frequency > 10^-5^). **g-h.** Fraction of variance in protein fitness explained by stability changes in domains grouped by the mode of inheritance of their associated disorders (g), or their disease mechanisms (h).

We next quantified the relationship between stability and pathogenicity in all individual domains with at least 20 annotated clinical variants (n = 17). Stability is the major contributor to pathogenicity in some domains. In the LIM domain 2 of FHL1, stability is an excellent classifier of pathogenic variants that cause reducing body myopathy (Matthew’s correlation coefficient, MCC = 1, Fig. 5c), a disease caused by the accumulation of FHL1 aggregates^38^ Similarly, the dominant Ankyloblepharon-Ectodermal Defects-Clefting (AEC) Syndrome is caused by mutations in the SAM domain of TP63 that lead to TP63 aggregation^39^. Accordingly, most pathogenic variants in the SAM domain of TP63 are destabilizing (MCC = 0.83, Fig. 5c).

In contrast, stability changes are a poorer predictor of pathogenic variants in other domains. A large proportion of mutations in the methyl-binding domain (MBD) of MECP2 that cause the dominant Rett Syndrome are not destabilizing (MCC = 0.4, Fig. 5c). This suggests that many of these haploinsufficient variants interfere with the methylated DNA binding function of MECP2^40–42^ without impacting the overall stability of the domain. Indeed, pathogenic mutations in MECP2 MBD that are not strongly destabilizing are concentrated in its DNA binding interface (OR = 5.00, p=0.09, Fig. 5d), in second-shell (OR = 4.09, p = 0.15) and in positively charged surface residues (OR = 4.35, p = 0.019), likely leading to a loss of binding affinity. Similarly, multiple mutations in the CRX homeodomain that cause inherited retinal dystrophies are not strongly destabilizing (MCC = 0.52, Fig. 5c). These variants are also enriched in its DNA binding interface (OR=6.33, p = 0.14 Fig. 5d) and in positively charged sites (OR = 3.58, p = 0.11). Interestingly, the mode of inheritance of mutations in CRX^43^ correlates with their stability effects: while the recessive R90W is strongly destabilizing (Fig. 5c), the dominant K88N and E80A are stable (Fig. 5c), consistent with their described gain-of-function mechanisms.^43^ This again suggests that destabilization is the major disease mechanism for some proteins and diseases but much less important for others.

### Protein stability in recessive and dominant disorders

Comparing across all domains with at least 20 clinical variants, there is a striking correlation between how well stability explains pathogenicity and how well stability explains evolutionary fitness, as quantified by ESM1v (Pearson’s r = 0.81, Fig. 4d). We therefore used the correlation between stability and evolutionary fitness to rank all 108 domains with at least one known pathogenic variant (Extended Data Fig. 5f). Domains where stability is highly predictive of fitness include many PHD finger and PDZ domains (Extended Data Fig. 5f). In these domains we expect stability changes to be the major driver of pathogenicity. In contrast, stability only poorly predicts the fitness of homeodomains, HMG-box, and nuclear hormone receptor-type zinc finger domains (Extended Data Fig. 5f), suggesting that other molecular mechanisms will more frequently cause pathogenicity.

The contribution of stability to protein fitness also varies among genes with different modes of inheritance and disease mechanisms (Fig. 5g,h). Recessive diseases are strongly associated with loss-of-function (LOF), while dominant diseases can also be caused by additional mechanisms such as gain-of-function and dominant negative effects, or toxic aggregation^45,46^. A median 44% of the variance in protein fitness is accounted for by stability changes in proteins mutated in recessive diseases, in contrast to only 26% in dominant disorders (p = 1.1 x 10^-3^, Wilcoxon rank sum test), suggesting that protein variants more frequently affect biophysical properties other than stability in dominant disorders. Indeed, LOF and aggregation diseases are better explained by protein destabilization than “altered function” diseases associated with gain-of-function or dominant negative mechanisms (Fig. 5h). Despite this overall association, however, we also find that within LOF diseases the extent to which destabilization explains protein fitness is variable (Fig. 5h). In summary, mutagenesis of >500 domains suggests that stability changes are an important cause of pathogenicity but that this varies across proteins, with changes in stability particularly important in recessive diseases.

### Conservation of mutational effects in homologous proteins

An important goal of Domainome 1 is to quantify the extent to which mutational effects are conserved in structurally homologous proteins. Mutations in homologous sites might be expected to have very similar effects. However, mutations can also interact energetically, resulting in changes in mutational effects depending upon the sequence context, a phenomenon known as epistasis^44^. If epistasis is prevalent then mutational effects will be poorly conserved in divergent homologous proteins, limiting the extent to which experimental data from some proteins can be used to predict stability changes in homologs^45^. Domainome 1 contains saturation mutagenesis data for five or more domains for 26 different domain families, allowing us to quantify the importance of epistasis for protein stability in structurally diverse protein folds.

To quantify the conservation of mutational effects, we fitted a thermodynamic model based on the Boltzmann partition function to all of the data for a domain family (Fig. 6a). The model assumes that mutations cause the same change in folding energy (ΔΔG) in all homologous domains, and that the energetic effects of mutations combine additively with no specific epistasis (Fig. 6a,b). We first fitted this model to homeodomains, the most abundant domain family in the dataset, with 37 homologs. The model provides very good prediction of mutational effects across all 37 homeodomains (Pearson’s r = 0.78 by ten-fold cross validation, Fig. 6c,d). The inferred free energy changes can be considered homolog-averaged mutational effects, providing an energy model describing the entire family (Fig. 6e,f). A linear model provides similarly good performance but with biased prediction residuals (r = 0.78, Extended Data Fig. 6). We additionally evaluated the performance of the Boltzmann energy model by leaving out single homeodomains from the training dataset (n = 37 models) and found similarly good performance (median Pearson’s r = 0.74). Predictive performance was, as expected, better for domains with more similar sequence to the training dataset, but reasonable across a wide range of sequence divergence (Pearson’s r = 0.73, Fig. 6g).

**Figure 6:**
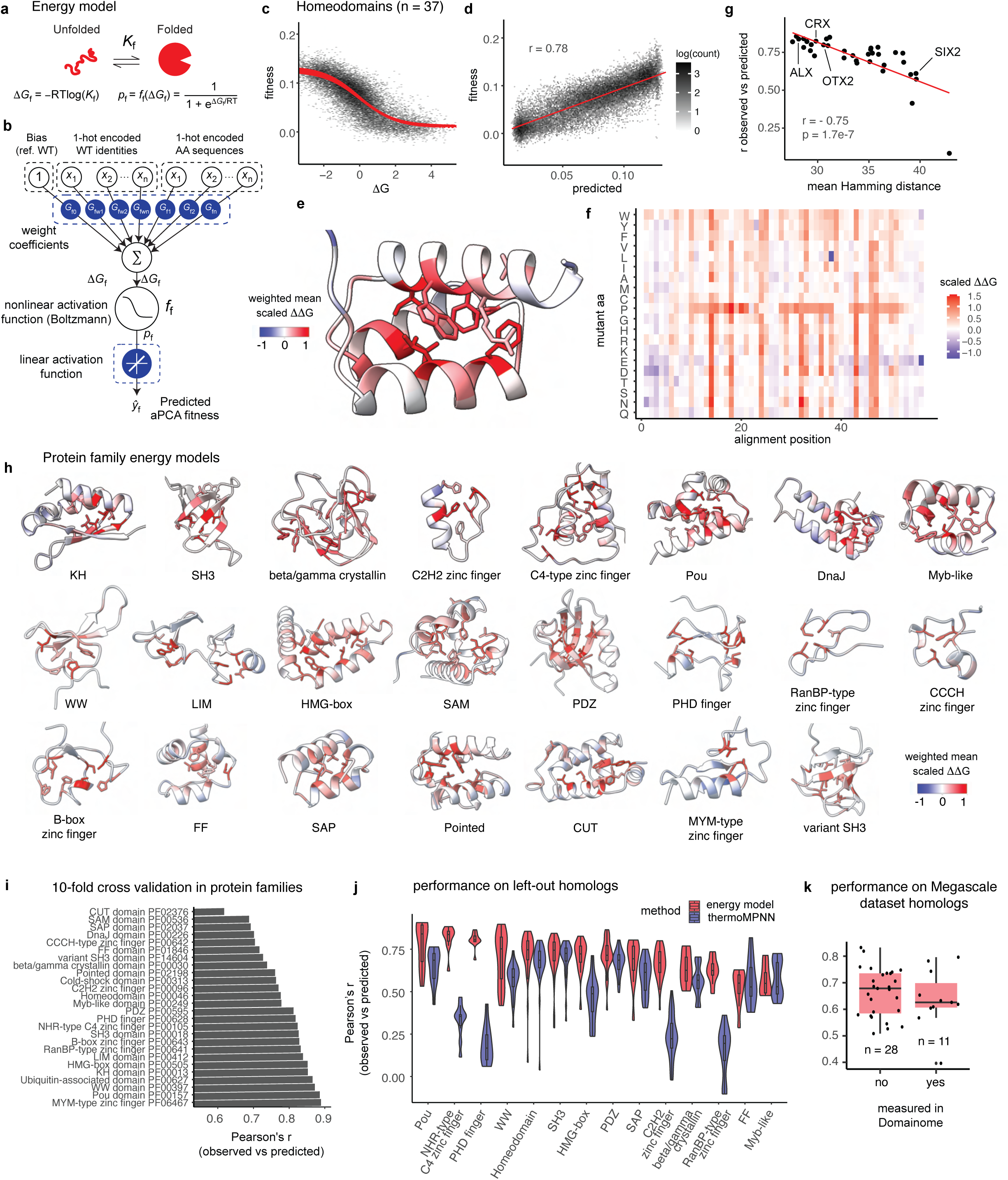
The genetic architecture of protein stability. **a.** Two-state folding equilibrium and corresponding thermodynamic model. ΔG_f_, Gibbs free energy of folding; K_f_, folding equilibrium constant; p_f_, fraction folded; ff, nonlinear function of ΔG_f_; R, gas constant; T, temperature in Kelvin. **b.** Neural network architecture used to fit thermodynamic models to protein families. **c.** Relationship between predicted fitness and additive trait (ΔG) in a Boltzmann model fit to the homeodomain family (PF00046). **d.** Correlation between observed fitness values and MoCHI fitness predictions of the homeodomain family for test set variants (10-fold cross validation; variants held out in any of the 10 training folds are shown). **e.** Structure of a representative homeodomain, with residues colored by the weighted mean homolog-averaged ΔΔG of mutations at each position. **f.** Heatmap depicting inferred homolog-averaged folding ΔΔG defining the stability of the homeodomain fold. **g.** Performance of the energy model on held out homeodomains as a function of the average genetic distance to the training set. **h.** Structures of representative domains of all modeled families with residues colored by the weighted mean homolog-averaged ΔΔG of mutations at each position. **i.** Summary of energy model performance on held out variants across protein families (10-fold cross validation). **j.** Summary of energy model performance on held out homologs across protein families, compared to the performance of the top stability predictor (ThermoMPNN). **k.** Summary of energy model performance on deep mutational scans of homologous domains generated using an *in vitro* proteolysis assay for stability determination^29^.

### The genetic architecture of protein stability across protein families

Extending this analysis to all 26 families with at least five homologs in Domainome 1, results in similarly accurate Boltzmann energy models for all families (median Pearson’s r = 0.80, median percent of explainable variance = 80.7%, Fig. 6i, Extended Data Fig. 7a,b). The performance on individual left out homologs is also very good (median Pearson’s r = 0.66, median percent of explainable variance = 73.5%, Fig. 6j). For most domains, predictions were best for the domains most similar to the training dataset but also good for domains with higher sequence divergence (Extended Data Fig. 7c). Predictions using these energy models were better than those made with ThermoMPNN, the top performing stability predictor on our dataset (Fig. 6j), and also had a good performance on stability deep mutagenesis scans generated using in vitro proteolysis selections^29^ (n = 38 homologs, median Pearson’s r = 0.65, Fig. 6k).

The excellent performance of these additive energy models is both useful and important: it demonstrates that epistasis makes only a small contribution to protein stability across these levels of sequence divergence. Combinatorial mutagenesis of individual proteins suggests a similar conclusion^46^. The decay of predictive performance with sequence divergence does, however, suggest a role for epistasis in the evolution of protein stability. Indeed, we identify 25,410 mutations with evidence of epistasis as variants with large residuals to the Boltzmann model fits (FDR < 0.1 Z-test, | residual | > 0.05 h^-1^, Extended Data Fig. 8a,b). These epistatic variants are enriched in the buried cores of protein domains (OR = 2.71, FET p = 1.05 x 10^-5^) and depleted from protein surfaces (OR = 0.19, FET p = 9.79 x 10^-14^, Extended Data Fig. 8c), across the full range of measured stabilities (Extended Data Fig. 8d) and protein families (Extended Data Fig. 8e). This data suggests that genetic interactions are more important for the evolution of protein cores, consistent with these residues having a larger number of structural contacts than solvent-exposed residues (Extended Data Fig. 8f).

### Energy models identify destabilizing mutations across entire domain families

Finally, the good performance of the Boltzmann energy models across all 26 families suggests that we can use them to provide proteome-wide stability predictions for these domain families. Using the models we made predictions for an additional 4,107,436 variants in 7,271 domains (Extended Data Fig. 9), including an additional 13,878 clinical variants, of which 1,310 are pathogenic, 951 are benign, and 11,617 are variants of uncertain significance (Fig. 7a,b). Consistent with the results for clinical variants with experimentally measured stability changes, 686 (52%) of these pathogenic variants are predicted to reduce protein stability (FDR < 0.1, OR = 2.95 compared to benign), and 452 (34%) are predicted to have strongly destabilizing effects (scaled ΔΔG > 0.3, FDR < 0.1, OR = 5.95 compared to benign, Fig. 7b,c). Similarly to the fitness-level data, our energy models outperform stability predictors in pathogenicity prediction, and have a lower performance than general variant effect predictors (Extended Data Fig. 9c). These high-quality stability predictions provide a resource for the mechanistic interpretation of clinical variants for entire protein families and suggest a strategy for expanding the Domainome proteome-wide by experimentally mutagenizing representative examples for all families.

**Figure 7:**
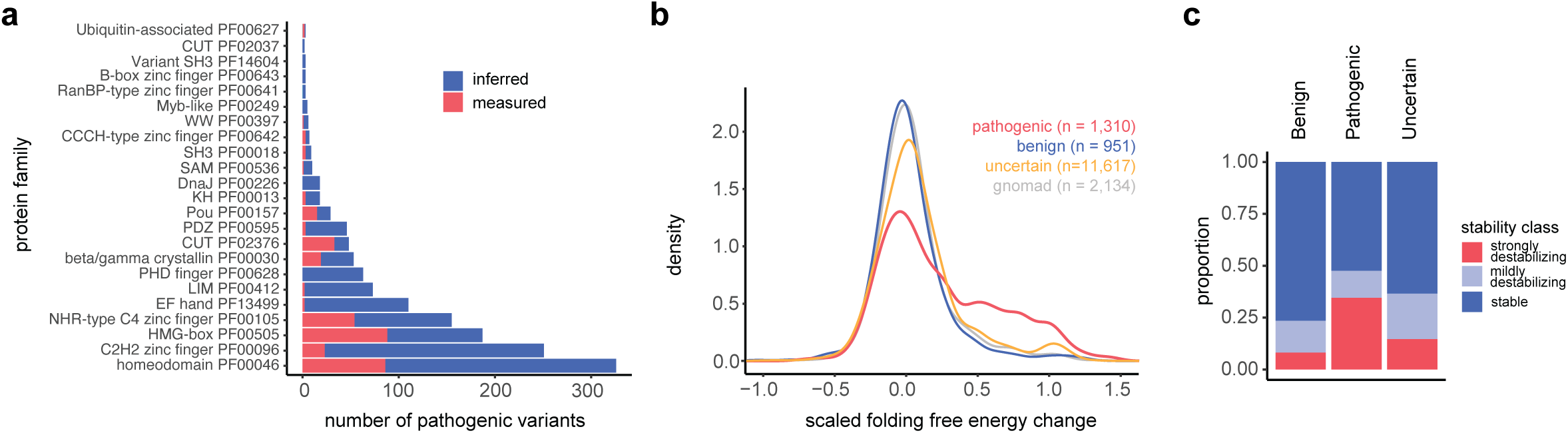
Extension of stability predictions to entire domain families. **a.** Number of pathogenic clinical variants covered by the predictions of homolog-averaged folding ΔΔGs. Colors represent whether variants are present in the aPCA fitness dataset (red) or not (blue). **b.** Distributions of scaled folding ΔΔG values of pathogenic, benign, uncertain, and gnomAD variants (allele frequency > 10^-5^). **c.** Proportion of strongly destabilizing variants (FDR < 0.1, scaled folding ΔΔG > 0.3) across classes of clinical variants.

## Discussion

We have presented here the first large-scale experimental analysis of mutational effects in human proteins. Human Domainome 1 demonstrates the feasibility of quantifying human protein variant effects at scale and provides high confidence measurements of changes in protein abundance for 563,534 variants in 522 structurally diverse protein domains. The dataset increases the number of stability measurements for human variants about 4.5-fold and serves as a large and standardized reference dataset for the interpretation of clinical variants and for benchmarking and training computational methods.

An important goal of this project was to evaluate the contribution of protein stability to genetic disease and sequence evolution. We found that at least 60% of pathogenic missense variants reduce protein stability but that this varies quite extensively across domains and diseases, with stability changes particularly important in recessive disorders. Similarly, we found that the contribution of stability to evolutionary fitness varies across protein families, with a larger contribution in all-beta protein domains compared to other fold types. How this relates to protein evolvability and the importance of trade-offs between stability and other molecular functions will be interesting avenues for future research. A further goal of Domainome 1 was to quantify the conservation of mutational effects in structurally homologous proteins. Fitting additive energy models to the data for domain families revealed that mutational effects on stability are largely conserved in homologous domains, with a small contribution from epistasis that increases with sequence divergence. This energetic additivity allows proteome-wide prediction of stability changes for entire families and suggests an efficient strategy to complete a first draft of the Domainome by mutagenizing representative examples for every family.

To maximize diversity and experimental efficiency, Domainome 1 focussed on structurally diverse small protein domains. An important caveat of this approach is the use of domains isolated from their native sequence context, and the extent to which mutational effects differ in full-length proteins is a key question for future work. Advances in DNA synthesis^47^, assembly^48^ and mutagenesis^49,50^ should facilitate this. Moreover, many domains did not yield sufficient signal in aPCA, indicating low stability or solubility, and other domains had mutational effects incompatible with two-state folding. In future work it will be important to understand the behavior of these domains in the assay and to test methods to increase the signal-to-noise such as altering domain boundaries, expression as full-length proteins, mutagenesis of exposed hydrophobic residues, and the use of different reporters or solubilization tags. In this study we have reported the effects of Domainome 1 variants on cellular protein abundance. However the same libraries can be re-used in future work to quantify variant effects on additional molecular traits, for example abundance in different cellular contexts^8,51,52^, *in vitro* fold stability^29^, protein-protein^53,54^ and protein-nucleic acid^55^ interactions, protein localisation^56^, aggregation^57^, and allostery^31,58^. It will also be important to include more complex genetic variants such as insertions and deletions^59^ and to extend to proteome-wide coverage, including extracellular and transmembrane proteins^60^.

Domainome 1 is a first important step in the comprehensive experimental analysis of human protein variants. It forms part of an ongoing global effort to determine the consequences of every mutation in every human protein and to produce reference atlases for the mechanistic interpretation of clinical variants^7^. Beyond human genetics, it is also part of a broader effort to produce large, well-calibrated datasets that quantify how changes in sequence alter the biophysical properties of proteins. We believe that such multimodal biophysical measurements for millions of proteins and variants will enable machine learning approaches to be effectively brought to bear on the generative functions of molecular biology, allowing accurate prediction and engineering from sequence^61^.

## Extended Data Figure legends

**Extended Data Figure 1: Mutating the human Domainome at scale.**

**a.** Histograms depicting sequencing coverage of the 9 site saturation mutagenesis libraries selected in this study. **b.** Correlations (Pearson’s r) between replicates for the 9 libraries. **c.** Distributions of quality control metrics in folded domains that were retained for further analyses, in folded domains excluded from further analysis, and in domains with less than 10% core residues or disordered. **d.** Correlations (Spearman’s rho) between aPCA scores and in vitro

ΔΔGs for individual protein domains. **e.** Correlations (Spearman’s rho) between aPCA scores and ΔΔGs derived through a high-throughput proteolysis assay for individual protein domains. **f.** Distributions of stability effects of mutations in protein cores and surfaces across the entire Domainome 1. Colors represent the physicochemical properties of the mutated amino acids (polar/charged = D, E, K, N, Q, R, S, T, hydrophobic = A, F, I, L, M, V, W, Y). **g.** Distributions of stability effects of mutations in different secondary structure types.

**Extended Data Figure 2: Deep mutational scans of 522 protein domains.**

Domains are ranked according to data quality (see Methods).

**Extended Data Figure 3: The contribution of protein stability to evolutionary constraint.**

**a.** Comparison of performance of variant effect and protein stability predictors in non-zinc finger domains (left) and zinc finger domains (right). The distribution of Spearman’s rho across all domains is shown for each predictor. **b.** Comparison of performance of variant effect and protein stability predictors in non-zinc finger domains with (right) and without (left) homology to protein domains in the Megascale dataset used for ThermoMPNN training. (left). **c.** Correlation between Pearson’s r and Spearman’s rho-based variance explained estimates of the contribution of stability to protein evolutionary constraint. **d.** Correlation between the contribution of stability to protein fitness estimated using aPCA data and using ThermoMPNN^34^ predictions, for non-zinc finger domains containing at least one pathogenic variant.

**Extended Data Figure 4: Identifying evolutionary functional sites using language models and abundance measurements.**

**a.** Representative examples of linear model and sigmoid model fits to the relationship between ESM1v predictions and abundance. **b.** Residuals to the fit of protein domain variants in several classes of annotated functional sites. Classes of functional sites with at least 50 residues annotated across the full domainome dataset are shown. **c.** Distribution of abundance to ESM1v residuals in functional sites, second shell sites, and other sites, for domains with at least 1 annotated functional site in CDD.

**Extended Data Figure 5: The contribution of protein destabilization to genetic disease.**

**a.** Gini plot depicting the cumulative number of pathogenic variants present in Domainome 1 domains ranked by pathogenic variant counts (from highest to lowest). **b.** Proportions of non-destabilizing, mildly destabilizing (FDR < 0.1, 0 > normalized aPCA fitness > -0.3), and strongly destabilizing (FDR < 0.1, normalized aPCA fitness < -0.3) variants in the pathogenic and benign sets, for several protein families. gnomAD variants with an allele frequency (af) > 1 x 10^-5^ were included as benign. Families with at least 10 variants in the pathogenic and in the benign classes are shown. **c.** ROC curves comparing clinical variant classification performance of aPCA data and variant effect or stability predictors. **d.** ROC curves comparing clinical variant classification performance of aPCA, ESM1v, protein structural features (secondary structure, rSASA, wild-type aa, mutant aa), and combinations of these using logistic regression (see Methods). **e.** Distributions of the proportions of variance in fitness explained by stability changes corrected by protein composition differences across domains. Protein domains are split according to the mode of inheritance of pathogenic mutations (left) or their disease mechanisms (right). **f.** Ranking of all 108 domains containing at least 1 pathogenic variant by the percentage of variance in fitness explained by stability changes (top left domains = highly explained by stability, bottom right domains = poorly explained by stability).

**Extended Data Figure 6: Comparison of linear and Boltzmann energy model fits to a protein family.**

Correlations between observed fitness values and energy model predictions for test set variants held out in any of the 10 training folds, for an energy model with a linear activation function (a) or a Boltzmann activation function (b) at the additive trait layer.

**Extended Data Figure 7: Thermodynamic models of protein families.**

**a.** Correlations between observed fitness values and energy model fitness predictions for test set variants held out in any of the 10 training folds, for Boltzmann models across protein families. **b.** Inferred homolog-averaged folding ΔΔGs across protein families, normalized such that the 2.5th percentile of the data corresponds to a scaled ΔΔG of -1. **c.** Correlation between model performance on left out protein domains, and the averaged genetic distance of the left-out domains to the rest of the training set, quantified as the hamming distance.

**Extended Data Figure 8: Protein cores are more epistatic than surfaces.**

**a-b.** Representative examples of an epistatic site (a) and a non-epistatic site (b) in the PDZ family. Epistatic mutations (FDR < 0.1, | residual | > 0.05 h^-1^) are shown in blue, and non-epistatic mutations are shown in pink. The epistatic site is enriched in epistatic mutations. **c.** Distributions of enrichment of epistatic mutations in protein core sites (rSASA<25% in >75% of homologs), surface sites (rSASA>25% in >75% of homologs), or changing sites. **d.** Enrichment of epistatic mutations in core (n = 368), surface (n = 1,039), and changing (n = 193) sites as a function of the average fitness of all variants per site. The running median of each class of sites across the fitness range (window size = 0.05 h^-1^) is shown as a colored line. **e.** Distribution of enrichment of epistatic mutations in core, surface, and changing sites across protein families. **f.** Distribution of total contact numbers per residue in core (n = 87,316) and surface (n = 193,797) residues across the Domainome, estimated from getcontacts.

**Extended Data Figure 9: Proteome-wide extension of stability predictions.**

**a.** Number of measured and predicted protein variants, by family. **b.** Number of measured and predicted protein domains, by family. **c.** ROC curves comparing clinical variant classification performance of ESM1v, FoldX, and energy models of protein families.

## Data availability

All DNA sequencing data have been deposited in the Gene Expression Omnibus under the accession number GSE265942. All aPCA measurements and their associated errors are available as Extended Data Table 2, including quality ranking by domain for data filtering. Weighted mean residuals of the comparisons between abundance and evolutionary fitness predictions are available as Extended Data Table 3. Homolog-averaged ΔΔG values and their associated errors mapped to homologous domains proteome-wide are available as Extended Data Table 4. aPCA scores and matching variant effect predictions are available as Extended Data Table 5.

## Code availability

Source code used to perform all analyses and to reproduce all figures in this work is available at: https://github.com/lehner-lab/domainome. Files required to reproduce the analyses can be downloaded at https://zenodo.org/records/11043643.

## Author contributions

A.B. designed the libraries and performed all experiments and analyses. X.J. and Y.S. synthesized the variant libraries. A.B. and B.L. conceived the project, designed analyses, and wrote the manuscript, with input from all authors.

## Acknowledgements

A.B. and B.L. were funded by a European Research Council (ERC) Advanced (883742) grant, the Spanish Ministry of Science and Innovation (LCF/PR/HR21/52410004, EMBL Partnership, Severo Ochoa Centre of Excellence), the Bettencourt Schueller Foundation, the AXA Research Fund, Agència de Gestió d’Ajuts Universitaris i de Recerca (AGAUR, 2017 SGR 1322), and the CERCA Program/Generalitat de Catalunya. A.B was funded by an EMBO (ALTF 183-2020) and a Marie Skłodowska-Curie (101030961) fellowship. X.J. and Y.S were funded by the National Natural Science Foundation of China (No.32322047) and Jiangsu Provincial Department of Science and Technology (No.BM2023009). We thank all members of the Lehner Lab for helpful discussions and suggestions.

## Materials and methods

### Media

- LB: 10 g/L Bacto-tryptone, 5 g/L Yeast extract, 10 g/L NaCl. Autoclaved 20 min at 120°C.
- YPDA: 20 g/L glucose, 20 g/L Peptone, 10 g/L Yeast extract, 40 mg/L adenine sulphate. Autoclaved 20 min at 120°C.
- SORB: 1 M sorbitol, 100 mM LiOAc, 10 mM Tris pH 8.0, 1 mM EDTA.
- Filter sterilized (0.2 mm Nylon membrane, ThermoScientific).
- Plate mixture: 40% PEG3350, 100 mM LiOAc, 10 mM Tris-HCl pH 8.0, 1 mM EDTA pH 8.0. Filter sterilized.
- Recovery medium: YPD (20 g/L glucose, 20 g/L Peptone, 10 g/L Yeast extract) +0.5 M sorbitol. Filter sterilized.
- SC -URA: 6.7 g/L Yeast Nitrogen base without amino acid, 20 g/L glucose, 0.77 g/L complete supplement mixture drop-out without uracil. Filter sterilized.
- SC -URA/ADE: 6.7 g/L Yeast Nitrogen base without amino acid, 20 g/L glucose, 0.76 g/L complete supplement mixture drop-out without uracil, adenine and methionine. Filter sterilized.
- MTX competition medium: SD –URA/ADE + 200 μg/mL methotrexate (BioShop Canada Inc., Canada), 2% DMSO.
- DNA extraction buffer: 2% Triton-X, 1% SDS, 100mM NaCl, 10mM Tris-HCl pH8, 1mM EDTA pH8.

### Library design, synthesis, and cloning

We sampled PFAM-annotated human protein domains (version 34.0) of intracellular human proteins (not defined as “extracellular”, “transmembrane”, or “secreted” in UniProt), prioritizing proteins that had at least one annotated pathogenic variant ClinVar^3^. We additionally included protein domains with *in vitro* stability measurements to benchmark the assay. We designed the library in two rounds: we first designed and selected two libraries (A1 and B3) containing single mutants of a total of 485 domains that had been previously tested to grow as WTsTo design second set of libraries (C1 to C7), based on the results of A1 and B3, we excluded domains without a well-defined hydrophobic core (not having at least 10% of residues with rSASA < 25%) and disordered domains defined as having an average AlphaFold2 pLDDT<50, indicative of protein disorder. This second set contained 631 domains that had been previously tested to grow as wildtype, and 132 additional domains not previously tested.

The domain sequences were codon optimized with emboss backtranseq^63^ using a *S. cerevisiae* codon usage table, and excluding the GCT alanine codon to prevent the appearance of HindIII restriction sites. The domain sequences were fused to 5% gggctgctctagaatggctagc and 3% aagcttggcggtggcgggtctg constant regions containing NheI and HindIII restriction sites for cloning. Libraries were synthesized by BGI using the mMPS technology (manuscript in preparation). The standard bases and degenerate bases N/K (representing the mixture of A, T, C and G, or the mixture of T and G respectively) were used for synthesis. The degenerated bases N/K are pre-mixed with an optimized ratio of each base to achieve an equal proportion of each amino acid at the mutation position. Compared with library A1 (sequence lengths ranging from 101 to 128 bases), all other eight libraries are relatively longer (sequence lengths ranging from 125 to 341 bases). To avoid the fact that the quality of synthesized DNA decreases when the length of DNA increases, we first segmented the sequences into 378,896 sub-sequences with length at ∼80-100 nt for synthesis using the mMPS synthesis system. Then we used polymerase cycling assembly (PCA)-PCR to construct each full-length DNA fragment of the libraries with the corresponding generated sub-fragments.

Libraries were cloned at a variant coverage of ∼100x or greater by restriction digestion and ligation into pGJJ162, an aPCA assay plasmid where the DHFR3 fragment is fused at the C-terminus of the target protein domains and the fusion is driven by the CYC promoter, and the DHFR1,2 fragment is expressed at high levels as is driven by the GPD promoter.

### Large-scale transformation and competition assay

Variant libraries were transformed in triplicate with a coverage of 20x or greater (with a mean coverage across libraries ∼170x). For each transformation, we grew a 1 L YPDA culture of late log phase *S. cerevisiae* BY4741 cells (OD_600_∼0.8-1), harvested the cells by centrifugation (5 minutes, 4000 g), resuspended in 43 mL SORB medium and incubated for 30 min on a shaker at room temperature. 875 μL 10 mg/mL previously boiled (5 min, 100 °C) ssDNA was added to the cells, followed by 17.5 μg library plasmid DNA and 175 mL Plate Mixture. The mix was incubated for 30 min on a shaker at room temperature. 17.5 mL DMSO were then added, and the cells were split in 50 mL Falcon tubes for 20 min heat shock at 42°C. Following incubation, cells were pooled and harvested by centrifugation, the supernatant was discarded with a pump, and cells were resuspended in 250 mL Recovery Media and incubated at 30°C for 1h. Cells were then centrifuged for 3 min at 3,000 g and transferred into 1 L SC –URA. 10 μL of this culture were immediately plated onto SC -URA selective plates to monitor transformation efficiency. The rest of the culture was incubated for one or two overnights at 30°C.

SC -URA cultures were used to inoculate a 1 L culture of SC -URA-ADE at an OD_600_=0.2-0.4, which was grown overnight (input culture). Cells from this culture were inoculated in 1 L SC -URA/ADE +200 μg/mL MTX to select stably expressed protein domain variants. The remaining input cells grown SC -URA/ADE were harvested and frozen for DNA extraction. MTX cultures were left to grow overnight to an OD_600_=1.6-2.5, corresponding approximately to 5 generations, harvested, and frozen for DNA extraction (outputs).

### DNA extraction, plasmid quantification and sequencing library preparation

Total DNA was extracted from yeast pellets equivalent to 50 mL of cells at OD_600_=1.6 as described in our previous work^31,58^. Plasmid concentrations in the resulting samples were quantified by against a standard curve of known concentrations by qPCR, using oGJJ152 and oGJJ153 as qPCR primers that amplify in the origin of replication of the aPCAa assay plasmid.

To generate the sequencing libraries, we performed two rounds of PCR amplification. In the first round, we used primer pools (oTB595+ and oTB748+) flanking the inserts that introduce frame-shifting nucleotides between the Illumina adapters and the sequencing region of interest. To maintain variant representation, we carried out eight 100 μL PCR1 reactions per sample, each of which starting with 125 million plasmid molecules that we amplified for 8 cycles. The reactions were column-purified (QIAquick PCR purification kit, QIAGEN), and the purified product was amplified further using the standard i5 and i7 primers to add the remainder of the Illumina adapter sequences and the demultiplexing indices (dual indexing) unique to each sample. We carried out a total of 8 100 μL PCR2 reactions per sample, each starting with 20-40 ng of purified product, that was amplified for 8 more cycles. The resulting amplicons were run on a 2% agarose gel to quantify and pool the samples for joint purification, and to ensure the specificity of the amplification and check for any potential excess amplification problems. The final libraries were size selected by electrophoresis on a 2% agarose gel, and gel-purified (QIAEX II Gel Extraction Kit, QIAGEN). The amplicons were subjected to Illumina paired end 2×150 sequencing on a NextSeq2000 instrument at the CRG Genomics facility.

### Sequencing data processing and normalization

FastQ files from paired-end sequencing of all aPCA selections were processed with DiMSum^64^ v1.2.11 (https://github.com/lehner-lab/DiMSum) using default settings with minor adjustments. The option “barcodeIdentityPath” was used to specify a variants file in order to recover designed variants only (NNK mutations in the protein domains present in each sublibrary), and starting and final culture optical densities and selection times were specified to infer absolute growth rates across libraries.

### Data filtering and normalization

We computed several quality control metrics on a per-domain basis. First, the wild-type position in the fitness distribution, as the difference between the wild-type and the 95th percentile of the distribution, divided by the difference between the 95th and 5th percentile in the distribution (fitness 90% range). Second, the Pearson’s correlation coefficient between replicates for missense variants. Third, the correlation between fitness and solvent accessibility for all variants in each domain. And fourth, the correlation between fitness and hydrophobicity of all variants in each domain (measured as the first principal component of a comprehensive table of amino acid properties^57^).

Compared to folded domains (more than 10% core residues and pLDDT>50), disordered and domains without a well-defined hydrophobic core (see *Library Design*) behave in a very distinct fashion in aPCA, with the wild type located in the middle of the fitness distribution, hydrophobicity negatively correlated with protein abundance, and narrow dynamic ranges resulting from typically small effects of mutations (Extended Data Fig. 1c). We thus excluded these domains from further analysis in the main text. Folded domains showed larger dynamic ranges, with the wild-type among the fittest variants, a negative correlation between mutation sensitivity and solvent accessibility, and a lack of correlation to hydrophobicity (Extended Data Fig 1c). For folded domains, the four quality metrics described above were highly correlated with each other, and we combined them using principal component analysis. The first principal component explained 64% of the variance, and was used as a single quality metric to rank all domains.

We retained all domains ranked 600 or higher, with a missense Pearson’s r > 0.485, and with more than 50% of variants measured (with at least 10 counts in at least 1 replicate), resulting in 538 domains. We removed from this set 16 additional domains that were not compatible with 2-state folding, either showing many large effect abundance-increasing mutations (O75364_PF00046_64, O75956_PF09806_73, P10242_PF00249_89, P52952_PF00046_140, Q13263_PF00643_205, Q86TZ1_PF13181_60, Q8IX03_PF00397_1, Q8NDW8_PF13181_799, Q9Y2H9_PF17820_968, Q9Y6M9_PF05347_15), narrow fitness ranges (P35637_PF00641_421, EHEE-rd2-0005_1, HHH-rd2-0133_1), or clear correlations to hydrophobicity (E9PAV3_PF19026_2040, HEEH-rd3-0223_1, Q5VTD9_PF00096_193), resulting in a final set of 522 domains.

The distributions of quality metrics for retained and discarded folded domains, and for disordered domains and folded domains without a well-defined core are in Extended Data Fig. 1c. Of the 478 discarded folded domains, 224 had slow growth rates as wt (< 0.075 h^-1^), 176 had an unfit wild type (wt position < 0.4), 107 were negatively correlated with hydrophobicity (Pearson’s r < -0.25), and 158 were not correlated with solvent accessibility (Pearson’s r < 0) (we note that many of the discarded domains meet more than one of these criteria). Discarded domains are shorter (median length = 49 aa) than retained domains (length = 58 aa, p = 1.87 x 10^-14^, Wilcoxon rank sum test), and are enriched in zinc-finger domains (57% of zinc finger domains discarded in contrast to 43%, 41%, and 39% for all-alpha, all-beta, and a+b, respectively, p = 2.33 x 10^-6^, Fisher’s exact test). Multimerization status only affected success rate marginally, as 49% of domains from proteins that form complexes^65^ were discarded, compared to 53% of those that do not engage in protein-protein interactions (p = 0.2, Fisher’s exact test). Finally, domains with yeast orthologs^66^ were less likely to be retained (45%) than domains without yeast orthologs (53%, p = 0.08, Fisher’s exact test).

We normalized the growth rates and growth rate errors within each protein domain by linearly scaling the data such that the WT normalized fitness equals zero, and the 2.5th percentile of the distribution of growth rates of all missense variants plus the WT is equivalent to a normalized fitness of -1. In all boxplots shown in the manuscript, the central line represents the median, the upper and lower hinges correspond to the first and third quartiles, with the upper whisker extending from the hinge to the largest value no further than 1.5 * IQR (inter-quartile range) from the hinge, and the lower whisker extending from the hinge to the smallest value no further than 1.5 * IQR from the hinge.

### Structural similarity network representation

We generated a structural distance matrix based on Foldseek^67^ hit probabilities. We computed all pairwise alignments between the domains in the final retained set by first creating a Foldseek database ‘foldseek_domainome_db’ containing all domains, and then searching all domains against the database using foldseek easy-search *.pdb foldseek_domainome_db foldseek_easy_allvsall tmp --format-output “query,target,alntmscore,qtmscore,ttmscore,prob” --exhaustive-search TRUE -e inf.

To visualize the similarity network, we imported the domains as nodes and the Foldseek probabilities as edges into Gephi^68^, and applied two layout algorithms: first the Fruchterman Reingold algorithm to equilibrium, which resulted in a clear separation of the different SCOP classes, followed by a Force Atlas 2 layout algorithm (preventing node overlap), which accentuated the separation between the different domain family clusters within each SCOP class.

### Comparison to reference ΔΔG datasets

To compare aPCA measurements with *in vitro* ΔΔGs, we used domains containing at least 10 variants in more than a single residue measured in vitro and in aPCA, and with a range of at least 2 kcal/mol for the in vitro measurements, and of 0.075 h^-1^ log growth rate units for the aPCA measurements (n = 10 domains). To compare aPCA measurements with ΔΔGs derived from high-throughput proteolysis deep mutational scans^29^, we used protein domains with an overlap (overlapping length / total alignment length) of at least 80%, and an aPCA measurement range of 0.075 h^-1^ log growth rate units (n = 12).

The total number of human missense variants measured in the mega-scale dataset^29^ was retrieved from the high-confidence dataset excluding double mutants and domain duplicates (from different PDB entries and genetic backgrounds of the same PDB entry). We also quantified the total number of human missense variants measured in all abundance (VAMP-seq, aPCA) datasets available (as of July 7th 2024) in MaveDB^62^ total number of human missense variants in ProthermDB^32^.

### Comparison to variant effect predictors

We generated ESM1v^12^ predictions (https://github.com/facebookresearch/esm) using the domain sequences alone as input, and using sliding windows across the human proteome (size=1000 aa, step=250 aa). For each domain, we used the predictions corresponding to the window in which the domain is most centered. We obtained EVE^11^, popEVE^69^ and Tranception^13^ scores from https://pop.evemodel.org/. ‘EVE domain’ scores computed on high-coverage alignments generated specifically for domainome domains were kindly shared by Aaron Kollasch and Debora Marks. We used precomputed RaSP scores^70^, and generated DDMut^71^ and ThermoMPNN^34^ predictions with the aid of AlphaFold2 structure predictions. FoldX^72^ predictions were obtained from https://ftp.ebi.ac.uk/pub/databases/ProtVar/predictions/stability/^65^. We used Spearman’s rho to quantify the relationship between the predictions and aPCA fitness. Domains with homology to protein domains in the Megascale dataset were defined using hmmer (http://hmmer.org/) hmmscan against PFAM, and predictors were evaluated on a homolog-free set to prevent leakage from ThermoMPNN training.

To estimate the fraction of the variance in mutational effects on evolutionary fitness that are attributable to protein stability, we calculated the correlation between ESM1v fitness predictions on full-length protein sequences and aPCA scores. The correlation coefficient was adjusted by the measurement error of the aPCA scores according to the Spearman disattenuation formula: 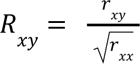 where *R*_*xy*_ is the disattenuated correlation coefficient, *R*_*xy*_ is the observed correlation coefficient, *r_xx_* is the mean correlation coefficient between aPCA replicates. This procedure was applied to both linear (Pearson’s) and rank (Spearman’s) correlation coefficients. The two were highly correlated (r = 0.94) and we report the Pearson’s r-based version for ease of interpretation.

### Analysis of functional sites

We carried out this analysis for domains where (1) the WT is above the 30th percentile in the fitness distribution, and (2) the range between the 5th and 95th percentile of the distribution of ESM1v predicted fitness is > 10 (n = 426 domains). We used sigmoid curves to model the relationship between normalized aPCA fitness (stability) and predicted function (ESM1v) of all variants in each individual domain, with an upper bound of 0 (WT aPCA normalized fitness) and a lower bound of -1:

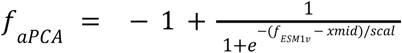

Where *f_aPCA_* is the aPCA normalized fitness, *f*_*ESM*1*v*_ is the ESM1v predicted fitness, *xmid* is the midpoint of the sigmoid and *scal* is the steepness parameter. To prioritize fitting the low stability variants, we weighted the fit by the aPCA normalized fitness:

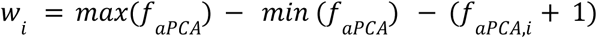

We used a two-tailed z-test to identify mutations whose effects on fitness cannot be accounted for by stability effects. We calculated z-scores as the aPCA residuals to the fit divided by the aPCA error, derived p-values based on the normal distribution, and performed multiple testing correction using Benjamini-Hochberg’s false discovery rate (FDR). We additionally calculated per-residue mean residuals weighted by the aPCA normalized fitness error.

We obtained residue-level functional site annotations corresponding to the Conserved Domains Database (CDD)^73^ using the InterPro API. Second-shell residues were defined as those with a minimum heavy atom distance of 5Å to functional site residues.

### Analysis of the stability effects of pathogenic variants

We identified destabilizing variants using a one-tailed z-test. Z-scores were calculated as the normalized fitness divided by the normalized fitness error, a p-value was derived on the basis of a normal distribution, and FDR multiple test correction was applied. Destabilizing variants were defined as variants with FDR < 0.1, and strongly destabilizing variants as FDR < 0.1 and a normalized aPCA fitness < -0.3. We obtained clinical variant annotations from ClinVar (January 2024 version) and from the UniProt API. Variant annotations were highly consistent between the two sources, and were merged. We tested for enrichments of clinical variant classes in stability classes using a two-tailed Fisher’s exact test.

To estimate to what extent destabilization explains pathogenic mutations in individual domains, we used Mathew’s correlation coefficient (MCC). To increase statistical power, we included gnomAD variants with an allele frequency > 10^-5^ as benign. We estimated the errors in MCCs by resampling the dataset based on the mean and errors of aPCA scores, 10 times. To analyze the distribution of pathogenic mutations in the MECP2 methyl-binding domain and the CRX homeodomain, we generated AlphaFold3 predictions^74^. DNA-binding interface residues were defined using getcontacts^75^, and second-shell residues were defined as those with a minimum heavy atom distance of 5Å to binding interface residues.

To extend the analysis to a larger number of domains with small numbers of pathogenic variants, we used the fraction of variance in ESM1v predicted fitness explained by aPCA scores as described above. MCCs based on clinical and gnomAD variants correlated well with the fraction of evolutionary fitness explained by stability effects in domains with at least 20 clinical and gnomAD variants, validating the approach. To further validate the estimates of the contribution of stability to fitness, we estimated the fraction of variance in ESM1v predicted fitness explained by thermoMPNN predicted stabilities for non-zinc finger domains with at least 1 pathogenic variant (n = 86). These correlated well with the original estimates derived using our abundance data (Pearson’s r = 0.81, Extended Data Figure 3d).

Modes of inheritance and mechanisms of disease information were obtained from OMIM (https://www.omim.org/). To analyze the contribution of stability changes to disease according to mode of inheritance and mechanism of action controlling for protein composition (ED Fig. 5e), we modeled the contribution of secondary structure and the percentage of core residues to the scores using linear models. We then extracted the model residuals as the composition-corrected scores.

### Clinical variant classification performance comparisons

We used the pROC R package^76^ to generate ROC curves and calculate ROC AUCs. We incorporated gnomAD variants with an allele frequency > 10^-5^ as benign. To test the performance when combining structural and sequence features (secondary structure, rSASA, wild-type residue, mutated residue), ESM1v, and aPCA scores, we used generalized linear models (glm) in R. In addition to training and evaluating on the full dataset, we trained the logistic regression models with 90% of the data and evaluated on the remaining 10% unseen data.

### Thermodynamic modeling of protein domain families

We used MoCHI^77^ to fit two-state thermodynamic models on a per-family basis (for families with at least 5 homologs and variants with a mean count >29). We specified a neural network architecture consisting of a single additive trait layer for shared folding energies across the family and a shared linear transformation layer. We used a “TwoStateFractionFolded” transformation derived from the Boltzmann distribution function that relates energies to proportions of folded protein. To map homologous positions across families, we used PFAM alignments. To input into MoCHI, we recoded each domain WT sequence as an indel sequence as long as the PFAM alignment, plus additional positions in the alignment with variants that encode the WT identity of each homolog. Mutations in each domain were encoded in the corresponding alignment position. This design allows MoCHI’s one-hot encoding of both the mutations and the WT identities simultaneously, and the joint fitting of mutation ΔΔG and starting ΔG of each homolog. We trained the model using a 10-fold cross-validation approach and evaluated the performance on held-out data.

We additionally fitted a linear model, and a “TwoStateFractionFolded” model with shared energies across all homologs of a family but with domain-specific linear transformations that account for potential differences in solubility of the folded and unfolded states between domains of the same family. These models resulted in highly correlated inferred energies to the original Boltzmann model (Pearson’s r = 0.972 for the linear model, and r = 0.938 for the Boltzmann model with domain-specific linear transformations) and similar performances on held out data across families. We discarded the linear model due to highly biased residuals, and chose the “TwoStateFractionFolded” model without homolog-specific linear scaling as the simplest of the remaining models. To compare across protein families, energies were rescaled such that the 2.5th percentile of the distribution of energies is equivalent to a scaled ΔΔG = -1 and the WT to a ΔΔG = 0.

We further evaluated the performance of the “TwoStateFractionFolded” model in families with at least 10 homologs by training the model leaving out a single domain at a time. The genetic distance (hamming distance and BLOSUM62 distance) of each left-out homolog to each of the homologs that went into training was calculated and averaged, to compare to model performance. The fraction of explainable variance in aPCA scores accounted for by the models was estimated as the R^2^ between observed and predicted fitness divided by the R^2^ between replicates.

### Epistasis analysis of protein domain families

We identified epistatic variants with large residuals to the MoCHI fits using a two-tailed z-test. We calculated z-scores as the residuals to the fit divided by the aPCA fitness errors, derived p-values on the basis of a normal distribution, and applied FDR multiple testing correction. Epistatic mutations were defined as those with |residual| > 0.05 and FDR < 0.1. Enrichments of alignment sites in epistatic variants were calculated, and significantly enriched sites were identified using a two-tailed Fisher’s exact test. Epistatic sites were defined as those with a log_2_(OR)>1.5 and a Fisher’s Exact test FDR < 0.05. We classified alignment sites as core residues (rSASA < 25% in at least 75% of homologs), surface residues (rSASA > 25% in at least 75% of homologs), or changing residues (the rest). We defined contacts using getcontacts^75^.

## Extended Data tables

**Extended Data table 1:** Library design: sequences, library statistics.

**Extended Data table 2:** Fitness scores and errors.

**Extended Data table 3:** Weighted mean residuals of abundance to evolutionary fitness predictions.

**Extended Data table 4:** Homolog-averaged ΔΔG predictions across families mapped to homologous domains proteome-wide.

**Extended Data table 5:** aPCA fitness scores and variant effect predictor scores.

**Extended Data table 6:**
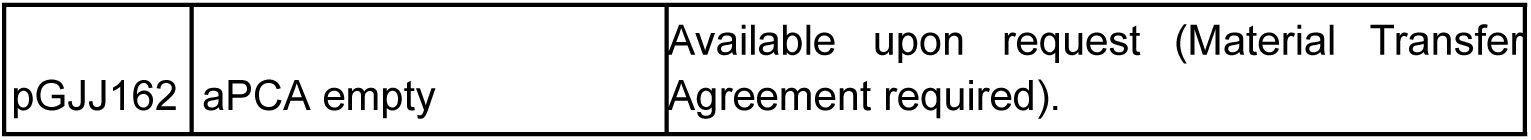
Plasmids.

**Extended Data table 7:**
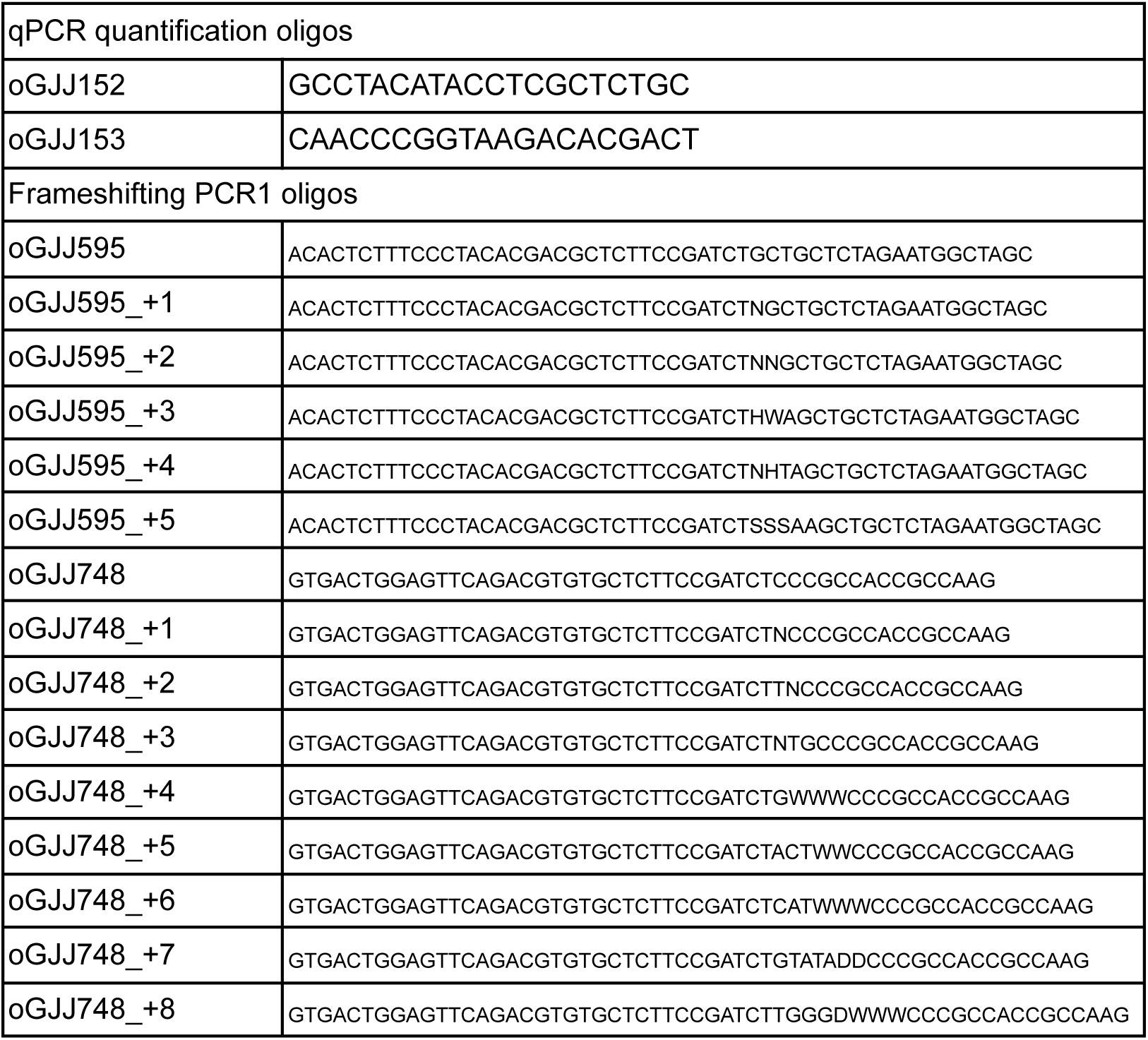
Oligonucleotides.

**Extended Data Table 8:** Sequences used in AF3 predictions MECP2 MBD bound to methylated DNA;

- MECP2 MBD: RGPMYDDPTLPEGWTRKLKQRKSGRSAGKYDVYLINPQGKAFRSKVELIAYFEKVGDT SLDPNDFDFTVTGR
- DNA chain 1: TCTGGAA-5mC-GGAATTCTTCTA
- DNA chain 2: TAGAAGAATTC-5meC-GTTCCAGA

CRX homeodomain bound to DNA:

- CRX homeodomain: RERTTFTRSQLEELEALFAKTQYPDVYAREEVALKINLPESRVQVWFKNRRAKCRQ
- DNA chain 1: ACGTGTGCACGTGATTAGTGCCATGCAACA
- DNA chain 2: TGTTGCATGGCACTAATCACGTGCACACGT

