## Extended data figures for "Site saturation mutagenesis of 500 human protein domains reveals the contribution of protein destabilization to genetic disease"

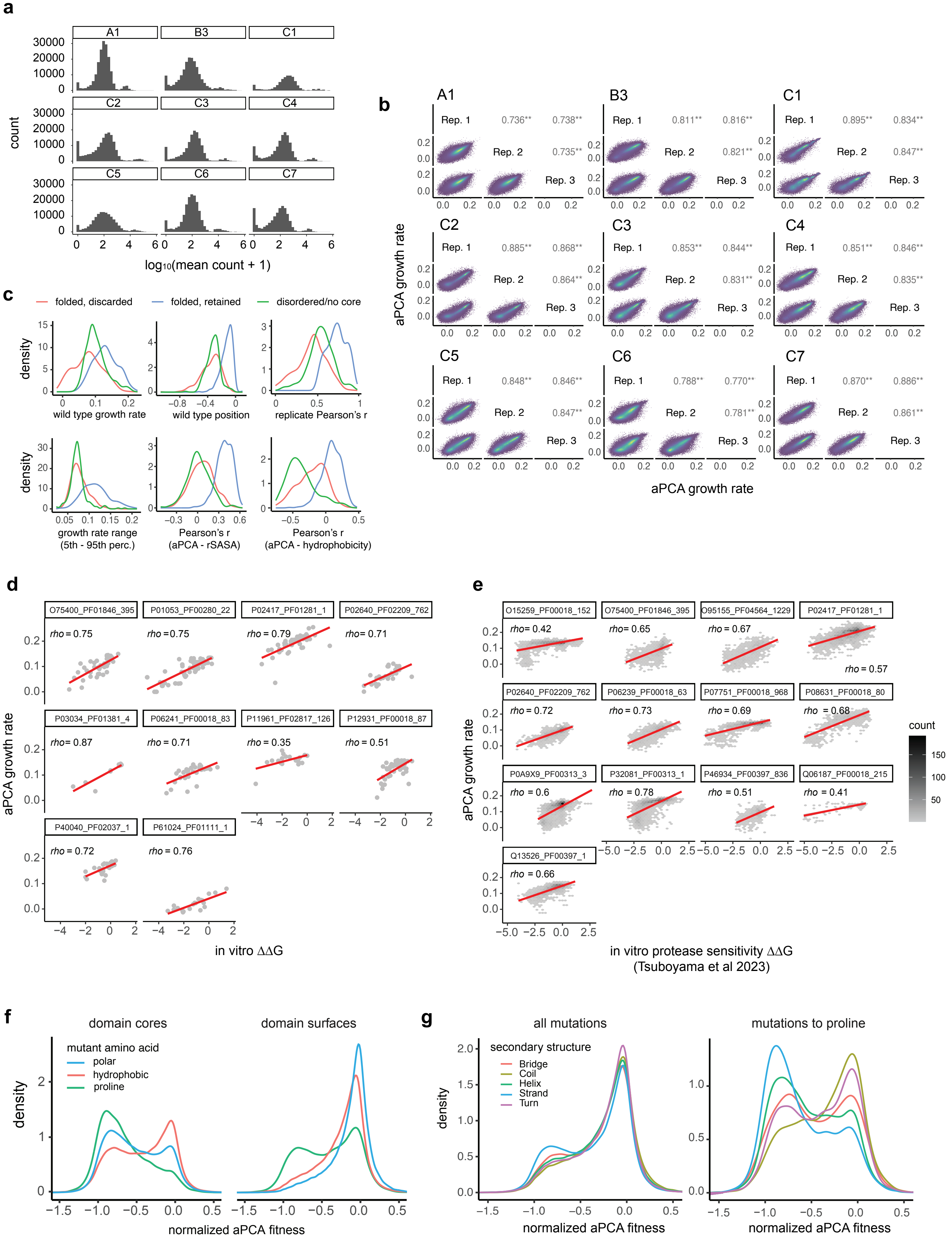

a
A1
B3
C1
C2
C3
C4
C5
C6
C7
count
log10(mean count + 1)
b
Rep. 1
Rep. 2
Rep. 3
aPCA growth rate
c
folded, discarded
folded, retained
disordered/no core
density
wild type growth rate
wild type position
replicate Pearson's r
growth rate range (5th - 95th perc.)
Pearson's r (aPCA - rSASA)
Pearson's r (aPCA - hydrophobicity)
d
aPCA growth rate
in vitro ΔΔG
e
count
in vitro protease sensitivity ΔΔG (Tsuboyama et al 2023)
f
domain cores
domain surfaces
mutant amino acid
polar
hydrophobic
proline
density
normalized aPCA fitness
g
all mutations
mutations to proline
secondary structure
Bridge
Coil
Helix
Strand
Turn
normalized aPCA fitness


ED Figure 1

### ED Figure 2-1

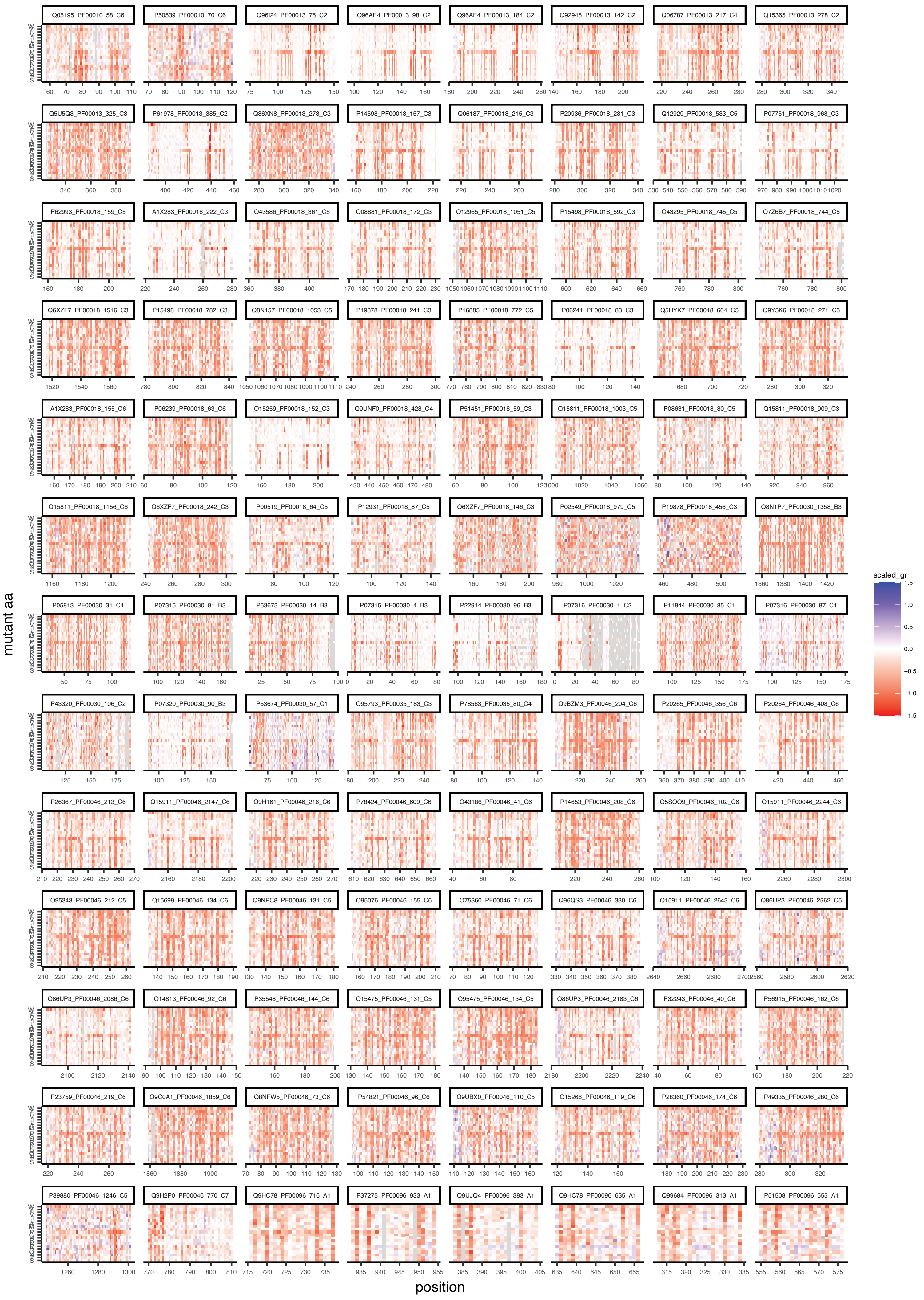

Q05195_PF00010_58_C6
P50539_PF00010_70_C6
Q96I24_PF00013_75_C2
Q96AE4_PF00013_98_C2
Q96AE4_PF00013_184_C2
Q92945_PF00013_142_C2
Q06787_PF00013_217_C4
Q15365_PF00013_278_C2
Q5U5Q3_PF00013_325_C3
P61978_PF00013_385_C2
Q86XN8_PF00013_273_C3
P14598_PF00018_157_C3
Q06187_PF00018_215_C3
P20936_PF00018_281_C3
Q12929_PF00018_533_C5
P07751_PF00018_968_C3
P62993_PF00018_159_C5
A1X283_PF00018_222_C3
O43586_PF00018_361_C5
Q08881_PF00018_172_C3
Q12965_PF00018_1051_C5
P15498_PF00018_592_C3
O43295_PF00018_745_C5
Q7Z6B7_PF00018_744_C5
Q6XZF7_PF00018_1516_C3
P15498_PF00018_782_C3
Q8N157_PF00018_1053_C5
P19878_PF00018_241_C3
P16885_PF00018_772_C5
P06241_PF00018_83_C3
Q5HYK7_PF00018_664_C5
Q9Y5K6_PF00018_271_C3
A1X283_PF00018_155_C6
P06239_PF00018_63_C6
O15259_PF00018_152_C3
Q9UNF0_PF00018_428_C4
P51451_PF00018_59_C3
Q15811_PF00018_1003_C5
P08631_PF00018_80_C5
Q15811_PF00018_909_C3
Q15811_PF00018_1156_C6
Q6XZF7_PF00018_242_C3
P00519_PF00018_64_C5
P12931_PF00018_87_C5
Q6XZF7_PF00018_146_C3
P02549_PF00018_979_C5
P19878_PF00018_456_C3
Q8N1P7_PF00030_1358_B3
P05813_PF00030_31_C1
P07315_PF00030_91_B3
P53673_PF00030_14_B3
P07315_PF00030_4_B3
P22914_PF00030_96_B3
P07316_PF00030_1_C2
P11844_PF00030_85_C1
P07316_PF00030_87_C1
P43320_PF00030_106_C2
P07320_PF00030_90_B3
P53674_PF00030_57_C1
O95793_PF00035_183_C3
P78563_PF00035_80_C4
Q9BZM3_PF00046_204_C6
P20265_PF00046_356_C6
P20264_PF00046_408_C6
P26367_PF00046_213_C6
Q15911_PF00046_2147_C6
Q9H161_PF00046_216_C6
P78424_PF00046_609_C6
O43186_PF00046_41_C6
P14653_PF00046_208_C6
Q5SQQ9_PF00046_102_C6
Q15911_PF00046_2244_C6
O95343_PF00046_212_C5
Q15699_PF00046_134_C6
Q9NPC8_PF00046_131_C5
O95076_PF00046_155_C6
O75360_PF00046_71_C6
Q96QS3_PF00046_330_C6
Q15911_PF00046_2643_C6
Q86UP3_PF00046_2562_C5
Q86UP3_PF00046_2086_C6
O14813_PF00046_92_C6
P35548_PF00046_144_C6
Q15475_PF00046_131_C5
O95475_PF00046_134_C5
Q86UP3_PF00046_2183_C6
P32243_PF00046_40_C6
P56915_PF00046_162_C6
P23759_PF00046_219_C6
Q9C0A1_PF00046_1859_C6
Q8NFW5_PF00046_73_C6
P54821_PF00046_96_C6
Q9UBX0_PF00046_110_C5
O15266_PF00046_119_C6
P28360_PF00046_174_C6
P49335_PF00046_280_C6
P39880_PF00046_1246_C5
Q9H2P0_PF00046_770_C7
Q9HC78_PF00096_716_A1
P37275_PF00096_933_A1
Q9UJQ4_PF00096_383_A1
Q9HC78_PF00096_635_A1
Q99684_PF00096_313_A1
P51508_PF00096_555_A1
mutant aa
position
scaled_gr
1.5
1.0
0.5
0.0
−0.5
−1.0
−1.5

ED Figure 2-2

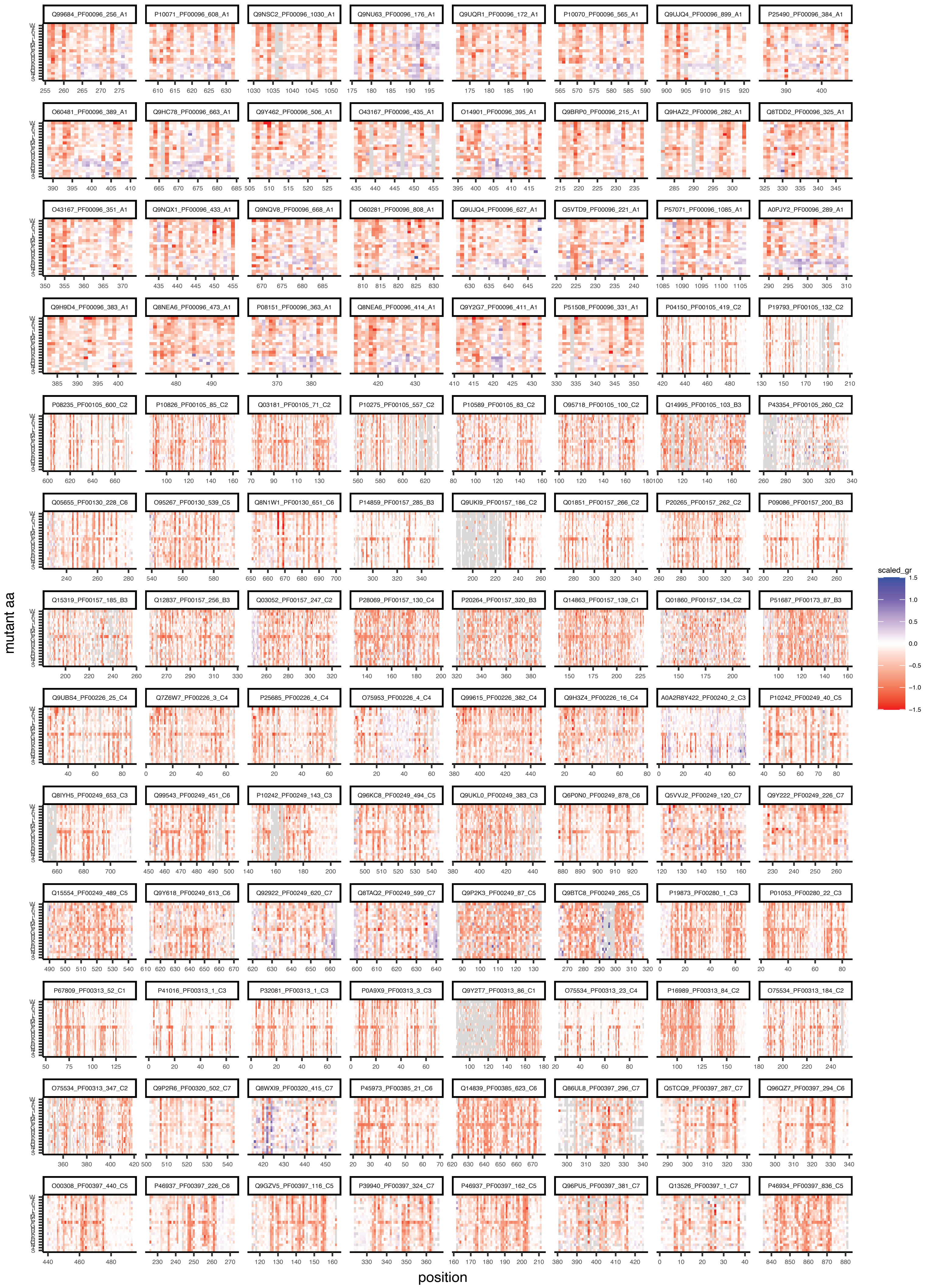

Q99684_PF00096_256_A1
P10071_PF00096_608_A1
Q9NSC2_PF00096_1030_A1
Q9NU63_PF00096_176_A1
Q9UQR1_PF00096_172_A1
P10070_PF00096_565_A1
Q9UJQ4_PF00096_899_A1
P25490_PF00096_384_A1
O60481_PF00096_389_A1
Q9HC78_PF00096_663_A1
Q9Y462_PF00096_506_A1
O43167_PF00096_435_A1
O14901_PF00096_395_A1
Q9BRP0_PF00096_215_A1
Q9HAZ2_PF00096_282_A1
Q8TDD2_PF00096_325_A1
O43167_PF00096_351_A1
Q9NQX1_PF00096_433_A1
Q9NQV8_PF00096_668_A1
O60281_PF00096_808_A1
Q9UJQ4_PF00096_627_A1
Q5VTD9_PF00096_221_A1
P57071_PF00096_1085_A1
A0PJY2_PF00096_289_A1
Q9H9D4_PF00096_383_A1
Q8NEA6_PF00096_473_A1
P08151_PF00096_363_A1
Q8NEA6_PF00096_414_A1
Q9Y2G7_PF00096_411_A1
P51508_PF00096_331_A1
P04150_PF00105_419_C2
P19793_PF00105_132_C2
P08235_PF00105_600_C2
P10826_PF00105_85_C2
Q03181_PF00105_71_C2
P10275_PF00105_557_C2
P10589_PF00105_83_C2
O95718_PF00105_100_C2
Q14995_PF00105_103_B3
P43354_PF00105_260_C2
Q05655_PF00130_228_C6
O95267_PF00130_539_C5
Q8N1W1_PF00130_651_C6
P14859_PF00157_285_B3
Q9UKI9_PF00157_186_C2
Q01851_PF00157_266_C2
P20265_PF00157_262_C2
P09086_PF00157_200_B3
Q15319_PF00157_185_B3
Q12837_PF00157_256_B3
Q03052_PF00157_247_C2
P28069_PF00157_130_C4
P20264_PF00157_320_B3
Q14863_PF00157_139_C1
Q01860_PF00157_134_C2
P51687_PF00173_87_B3
Q9UBS4_PF00226_25_C4
Q7Z6W7_PF00226_3_C4
P25685_PF00226_4_C4
O75953_PF00226_4_C4
Q99615_PF00226_382_C4
Q9H3Z4_PF00226_16_C4
A0A2R8Y422_PF00240_2_C3
P10242_PF00249_40_C5
Q8IYH5_PF00249_653_C3
Q99543_PF00249_451_C6
P10242_PF00249_143_C3
Q96KC8_PF00249_494_C5
Q9UKL0_PF00249_383_C3
Q6P0N0_PF00249_878_C6
Q5VVJ2_PF00249_120_C7
Q9Y222_PF00249_226_C7
Q15554_PF00249_489_C5
Q9Y618_PF00249_613_C6
Q92922_PF00249_620_C7
Q8TAQ2_PF00249_599_C7
Q9P2K3_PF00249_87_C5
Q9BTC8_PF00249_265_C5
P19873_PF00280_1_C3
P01053_PF00280_22_C3
P67809_PF00313_52_C1
P41016_PF00313_1_C3
P32081_PF00313_1_C3
P0A9X9_PF00313_3_C3
Q9Y2T7_PF00313_86_C1
O75534_PF00313_23_C4
P16989_PF00313_84_C2
O75534_PF00313_184_C2
O75534_PF00313_347_C2
Q9P2R6_PF00320_502_C7
Q8WXI9_PF00320_415_C7
P45973_PF00385_21_C6
Q14839_PF00385_623_C6
Q86UL8_PF00397_296_C7
Q5TCQ9_PF00397_287_C7
Q96QZ7_PF00397_294_C6
O00308_PF00397_440_C5
P46937_PF00397_226_C6
Q9GZV5_PF00397_116_C5
P39940_PF00397_324_C7
P46937_PF00397_162_C5
Q96PU5_PF00397_381_C7
Q13526_PF00397_1_C7
P46934_PF00397_836_C5
mutant aa
position
scaled_gr
1.5
1.0
0.5
0.0
−0.5
−1.0
−1.5

### ED Figure 2-3

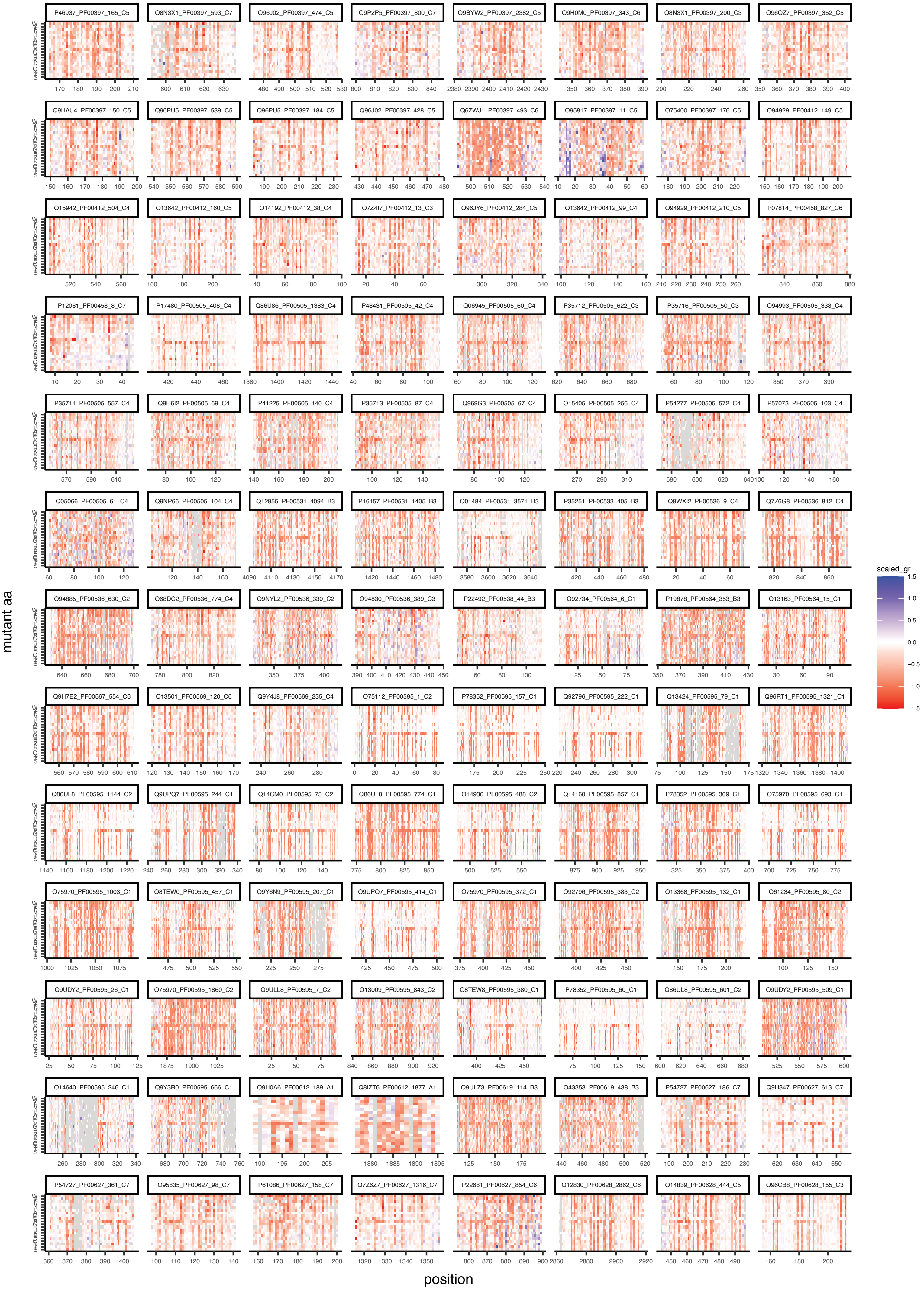

P46937_PF00397_165_C5
Q8N3X1_PF00397_593_C7
Q96J02_PF00397_474_C5
Q9P2P5_PF00397_800_C7
Q9BYW2_PF00397_2382_C5
Q9H0M0_PF00397_343_C6
Q8N3X1_PF00397_200_C3
Q96QZ7_PF00397_352_C5
Q9HAU4_PF00397_150_C5
Q96PU5_PF00397_539_C5
Q96PU5_PF00397_184_C5
Q96J02_PF00397_428_C5
Q6ZWJ1_PF00397_493_C6
O95817_PF00397_11_C5
O75400_PF00397_176_C5
O94929_PF00412_149_C5
Q15942_PF00412_504_C4
Q13642_PF00412_160_C5
Q14192_PF00412_38_C4
Q7Z4I7_PF00412_13_C3
Q96JY6_PF00412_284_C5
Q13642_PF00412_99_C4
O94929_PF00412_210_C5
P07814_PF00458_827_C6
P12081_PF00458_8_C7
P17480_PF00505_408_C4
Q86U86_PF00505_1383_C4
P48431_PF00505_42_C4
Q06945_PF00505_60_C4
P35712_PF00505_622_C3
P35716_PF00505_50_C3
O94993_PF00505_338_C4
P35711_PF00505_557_C4
Q9H6I2_PF00505_69_C4
P41225_PF00505_140_C4
P35713_PF00505_87_C4
Q969G3_PF00505_67_C4
O15405_PF00505_256_C4
P54277_PF00505_572_C4
P57073_PF00505_103_C4
Q05066_PF00505_61_C4
Q9NP66_PF00505_104_C4
Q12955_PF00531_4094_B3
P16157_PF00531_1405_B3
Q01484_PF00531_3571_B3
P35251_PF00533_405_B3
Q8WXI2_PF00536_9_C4
Q7Z6G8_PF00536_812_C4
O94885_PF00536_630_C2
Q68DC2_PF00536_774_C4
Q9NYL2_PF00536_330_C2
O94830_PF00536_389_C3
P22492_PF00538_44_B3
Q92734_PF00564_6_C1
P19878_PF00564_353_B3
Q13163_PF00564_15_C1
Q9H7E2_PF00567_554_C6
Q13501_PF00569_120_C6
Q9Y4J8_PF00569_235_C4
O75112_PF00595_1_C2
P78352_PF00595_157_C1
Q92796_PF00595_222_C1
Q13424_PF00595_79_C1
Q96RT1_PF00595_1321_C1
Q86UL8_PF00595_1144_C2
Q9UPQ7_PF00595_244_C1
Q14CM0_PF00595_75_C2
Q86UL8_PF00595_774_C1
O14936_PF00595_488_C2
Q14160_PF00595_857_C1
P78352_PF00595_309_C1
O75970_PF00595_693_C1
O75970_PF00595_1003_C1
Q8TEW0_PF00595_457_C1
Q9Y6N9_PF00595_207_C1
Q9UPQ7_PF00595_414_C1
O75970_PF00595_372_C1
Q92796_PF00595_383_C2
Q13368_PF00595_132_C1
Q61234_PF00595_80_C2
Q9UDY2_PF00595_26_C1
O75970_PF00595_1860_C2
Q9ULL8_PF00595_7_C2
Q13009_PF00595_843_C2
Q8TEW8_PF00595_380_C1
P78352_PF00595_60_C1
Q86UL8_PF00595_601_C2
Q9UDY2_PF00595_509_C1
O14640_PF00595_246_C1
Q9Y3R0_PF00595_666_C1
Q9H0A6_PF00612_189_A1
Q8IZT6_PF00612_1877_A1
Q9ULZ3_PF00619_114_B3
O43353_PF00619_438_B3
P54727_PF00627_186_C7
Q9H347_PF00627_613_C7
P54727_PF00627_361_C7
O95835_PF00627_98_C7
P61086_PF00627_158_C7
Q7Z6Z7_PF00627_1316_C7
P22681_PF00627_854_C6
Q12830_PF00628_2862_C6
Q14839_PF00628_444_C5
Q96CB8_PF00628_155_C3
scaled_gr
1.5
1.0
0.5
0.0
−0.5
−1.0
−1.5
mutant aa
position

ED Figure 2-4

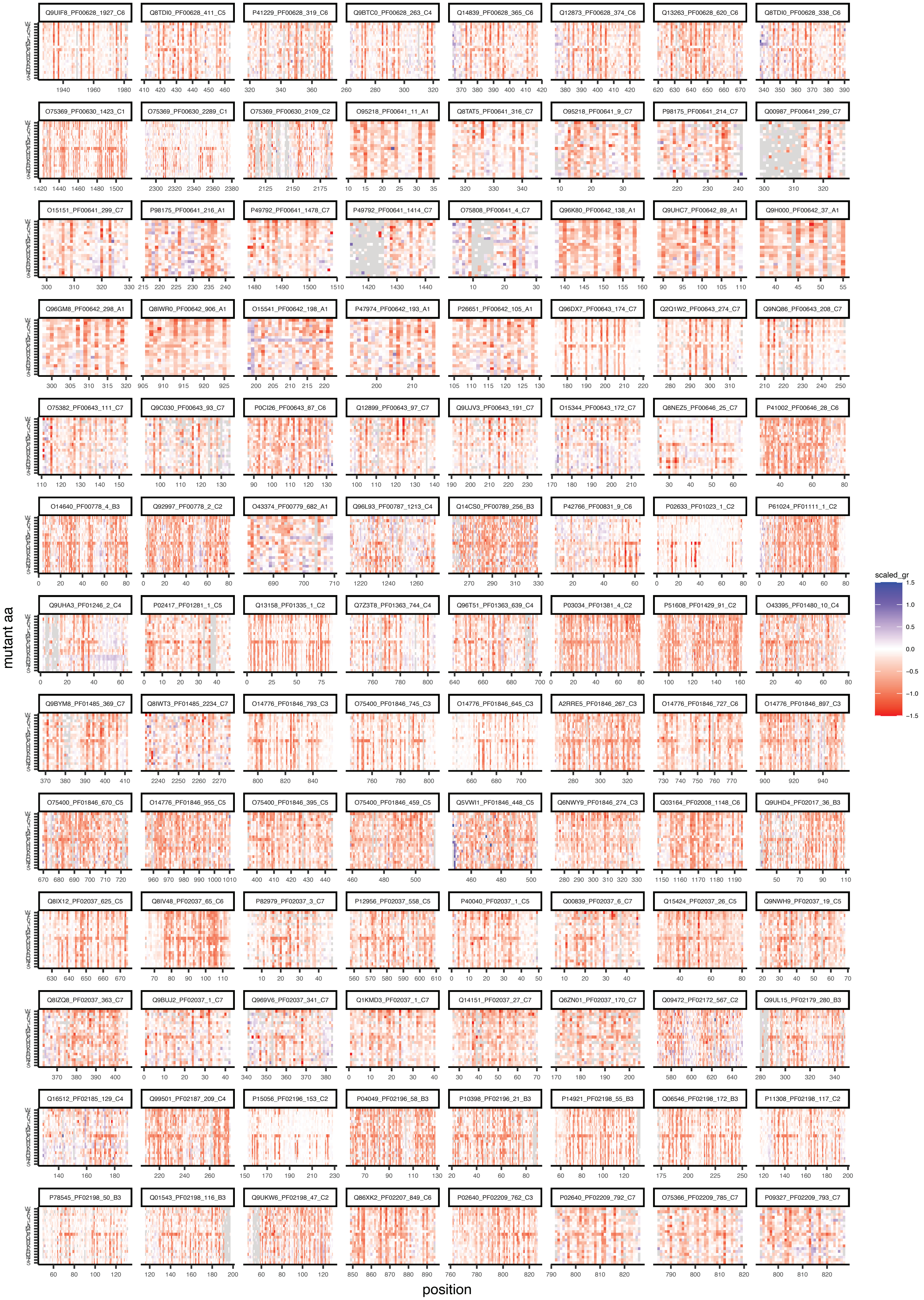

Q9UIF8_PF00628_1927_C6
Q8TDI0_PF00628_411_C5
P41229_PF00628_319_C6
Q9BTC0_PF00628_263_C4
Q14839_PF00628_365_C6
Q12873_PF00628_374_C6
Q13263_PF00628_620_C6
Q8TDI0_PF00628_338_C6
O75369_PF00630_1423_C1
O75369_PF00630_2289_C1
O75369_PF00630_2109_C2
O95218_PF00641_11_A1
Q8TAT5_PF00641_316_C7
O95218_PF00641_9_C7
P98175_PF00641_214_C7
Q00987_PF00641_299_C7
O15151_PF00641_299_C7
P98175_PF00641_216_A1
P49792_PF00641_1478_C7
P49792_PF00641_1414_C7
O75808_PF00641_4_C7
Q96K80_PF00642_138_A1
Q9UHC7_PF00642_89_A1
Q9H000_PF00642_37_A1
Q96GM8_PF00642_298_A1
Q8IWR0_PF00642_906_A1
O15541_PF00642_198_A1
P47974_PF00642_193_A1
P26651_PF00642_105_A1
Q96DX7_PF00643_174_C7
Q2Q1W2_PF00643_274_C7
Q9NQ86_PF00643_208_C7
O75382_PF00643_111_C7
Q9C030_PF00643_93_C7
P0CI26_PF00643_87_C6
Q12899_PF00643_97_C7
Q9UJV3_PF00643_191_C7
O15344_PF00643_172_C7
Q8NEZ5_PF00646_25_C7
P41002_PF00646_28_C6
O14640_PF00778_4_B3
Q92997_PF00778_2_C2
O43374_PF00779_682_A1
Q96L93_PF00787_1213_C4
Q14CS0_PF00789_256_B3
P42766_PF00831_9_C6
P02633_PF01023_1_C2
P61024_PF01111_1_C2
Q9UHA3_PF01246_2_C4
P02417_PF01281_1_C5
Q13158_PF01335_1_C2
Q7Z3T8_PF01363_744_C4
Q96T51_PF01363_639_C4
P03034_PF01381_4_C2
P51608_PF01429_91_C2
O43395_PF01480_10_C4
Q9BYM8_PF01485_369_C7
Q8IWT3_PF01485_2234_C7
O14776_PF01846_793_C3
O75400_PF01846_745_C3
O14776_PF01846_645_C3
A2RRE5_PF01846_267_C3
O14776_PF01846_727_C6
O14776_PF01846_897_C3
O75400_PF01846_670_C5
O14776_PF01846_955_C5
O75400_PF01846_395_C5
O75400_PF01846_459_C5
Q5VWI1_PF01846_448_C5
Q6NWY9_PF01846_274_C3
Q03164_PF02008_1148_C6
Q9UHD4_PF02017_36_B3
Q8IX12_PF02037_625_C5
Q8IV48_PF02037_65_C6
P82979_PF02037_3_C7
P12956_PF02037_558_C5
P40040_PF02037_1_C5
Q00839_PF02037_6_C7
Q15424_PF02037_26_C5
Q9NWH9_PF02037_19_C5
Q8IZQ8_PF02037_363_C7
Q9BUJ2_PF02037_1_C7
Q969V6_PF02037_341_C7
Q1KMD3_PF02037_1_C7
Q14151_PF02037_27_C7
Q6ZN01_PF02037_170_C7
Q09472_PF02172_567_C2
Q9UL15_PF02179_280_B3
Q16512_PF02185_129_C4
Q99501_PF02187_209_C4
P15056_PF02196_153_C2
P04049_PF02196_58_B3
P10398_PF02196_21_B3
P14921_PF02198_55_B3
Q06546_PF02198_172_B3
P11308_PF02198_117_C2
P78545_PF02198_50_B3
Q01543_PF02198_116_B3
Q9UKW6_PF02198_47_C2
Q86XK2_PF02207_849_C6
P02640_PF02209_762_C3
P02640_PF02209_792_C7
O75366_PF02209_785_C7
P09327_PF02209_793_C7
mutant aa
position
scaled_gr
1.5
1.0
0.5
0.0
−0.5
−1.0
−1.5

### ED Figure 2-5

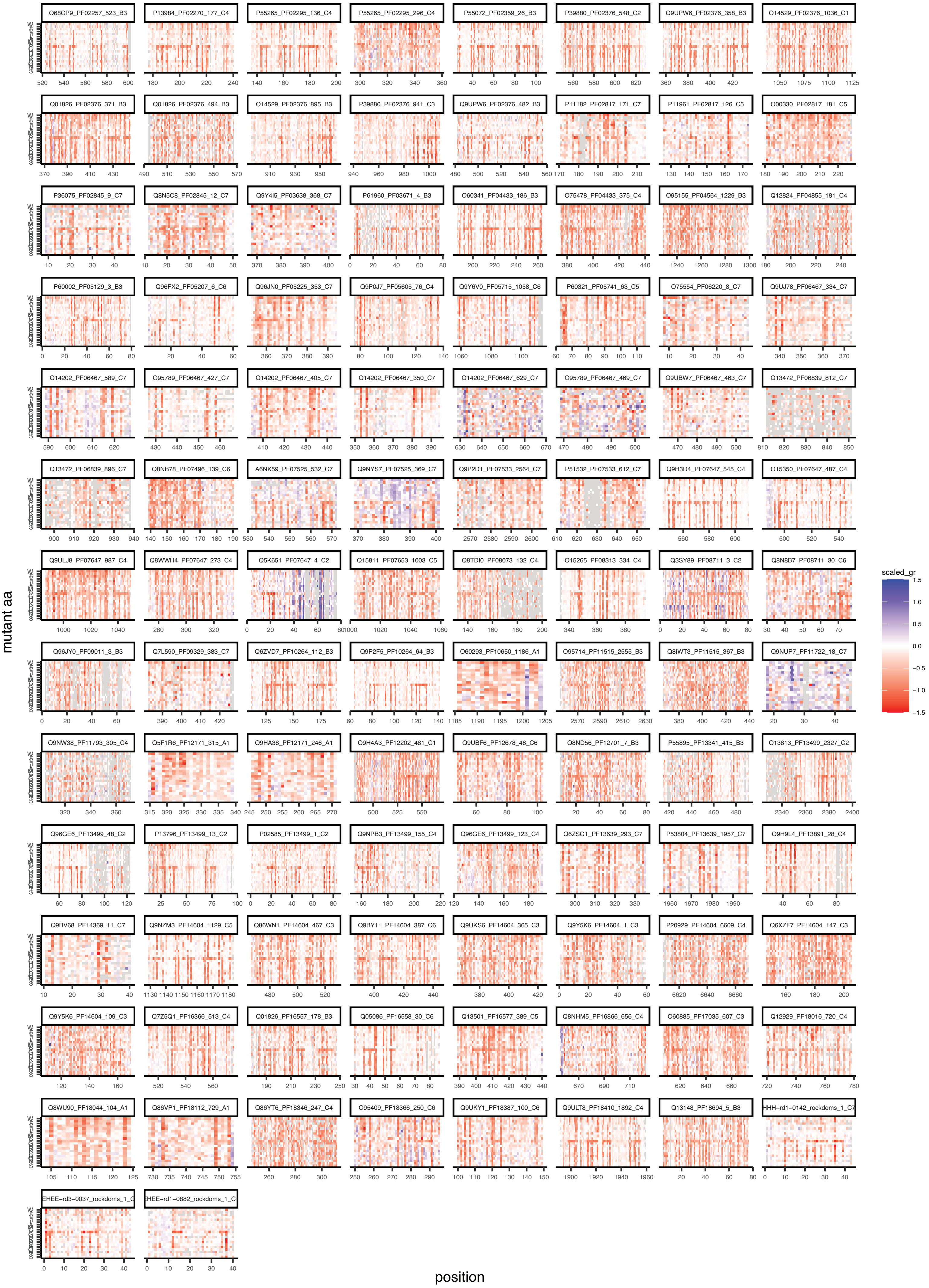

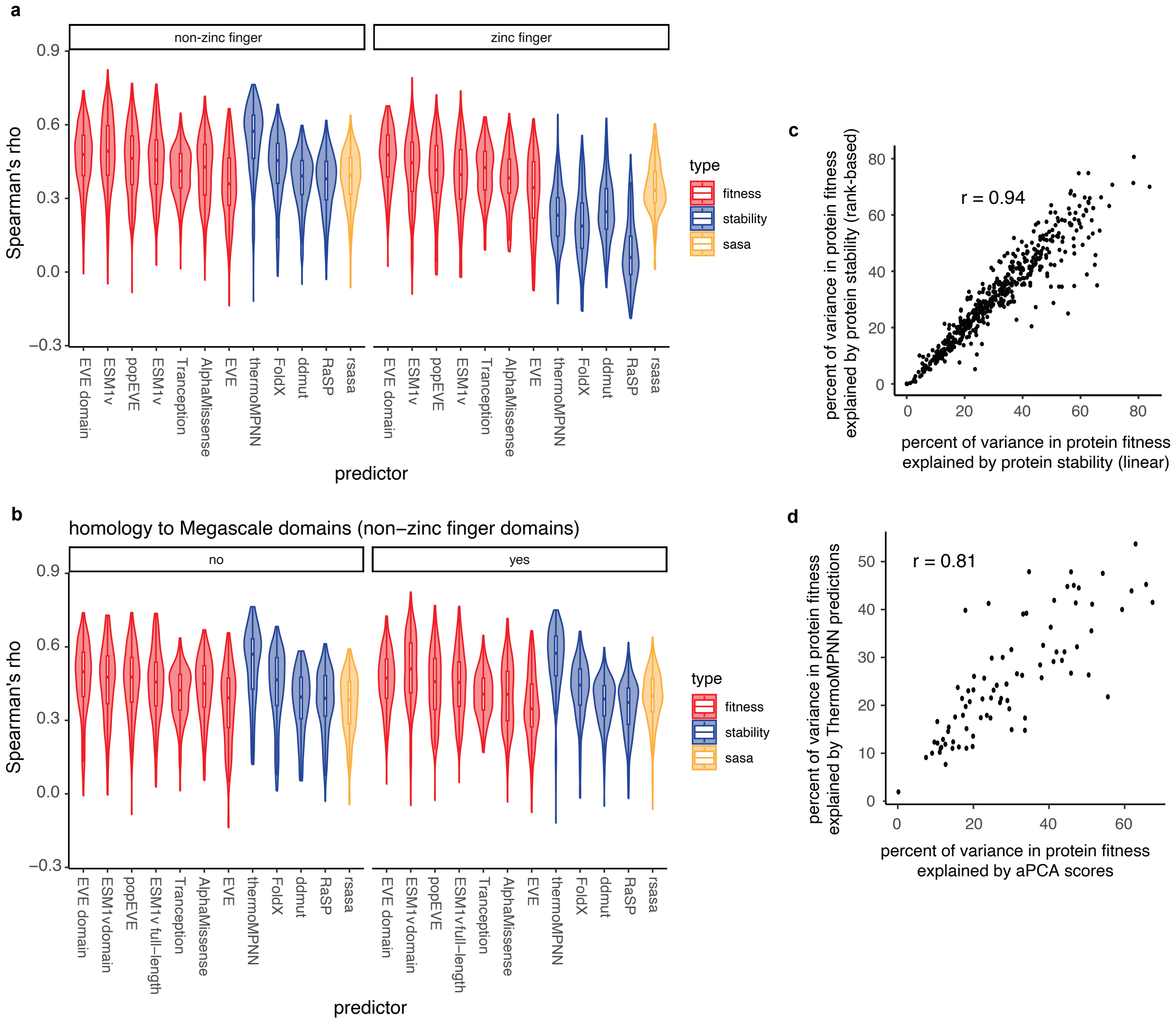
a
non-zinc finger
zinc finger
Spearman's rho
0.9
0.6
0.3
0.0
−0.3
EVE domain
ESM1v
popEVE
ESM1v
Tranception
AlphaMissense
EVE
thermoMPNN
FoldX
ddmut
RaSP
rsasa
predictor
type
fitness
stability
sasa
b
homology to Megascale domains (non−zinc finger domains)
no
yes
EVE domain
ESM1vdomain
popEVE
ESM1v full−length
Tranception
AlphaMissense
EVE
thermoMPNN
FoldX
ddmut
RaSP
rsasa
c
r = 0.94
percent of variance in protein fitness
explained by protein stability (rank-based)
percent of variance in protein fitness
explained by protein stability (linear)
d
r = 0.81
percent of variance in protein fitness
explained by ThermoMPNN predictions
percent of variance in protein fitness
explained by aPCA scores

ED Figure 3

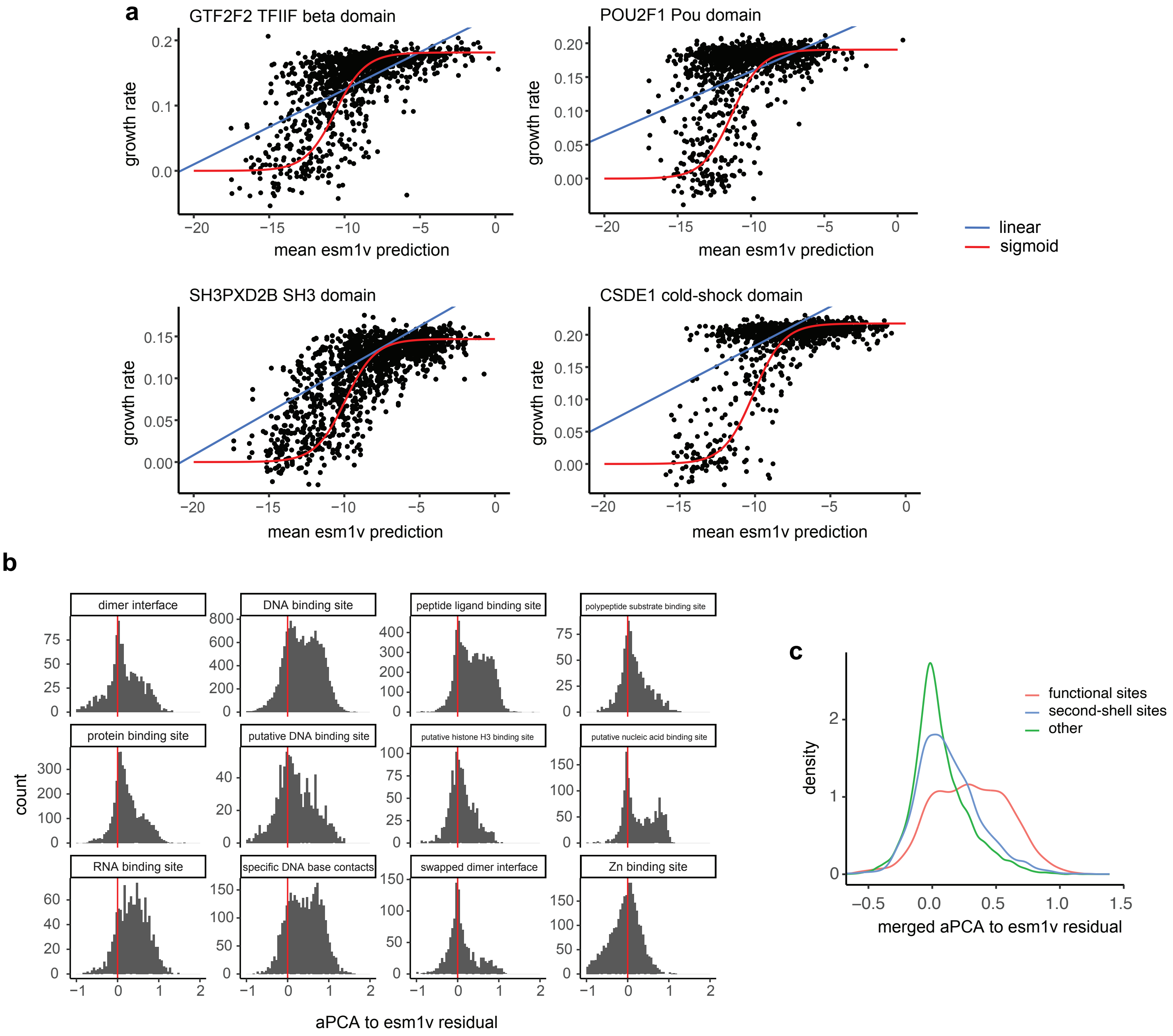
a
GTF2F2 TFIIF beta domain
POU2F1 Pou domain
SH3PXD2B SH3 domain
CSDE1 cold-shock domain
growth rate
mean esm1v prediction
linear
sigmoid
b
dimer interface
DNA binding site
peptide ligand binding site
polypeptide substrate binding site
protein binding site
putative DNA binding site
putative histone H3 binding site
putative nucleic acid binding site
RNA binding site
specific DNA base contacts
swapped dimer interface
Zn binding site
count
aPCA to esm1v residual
c
functional sites
second-shell sites
other
density
merged aPCA to esm1v residual

ED Figure 4

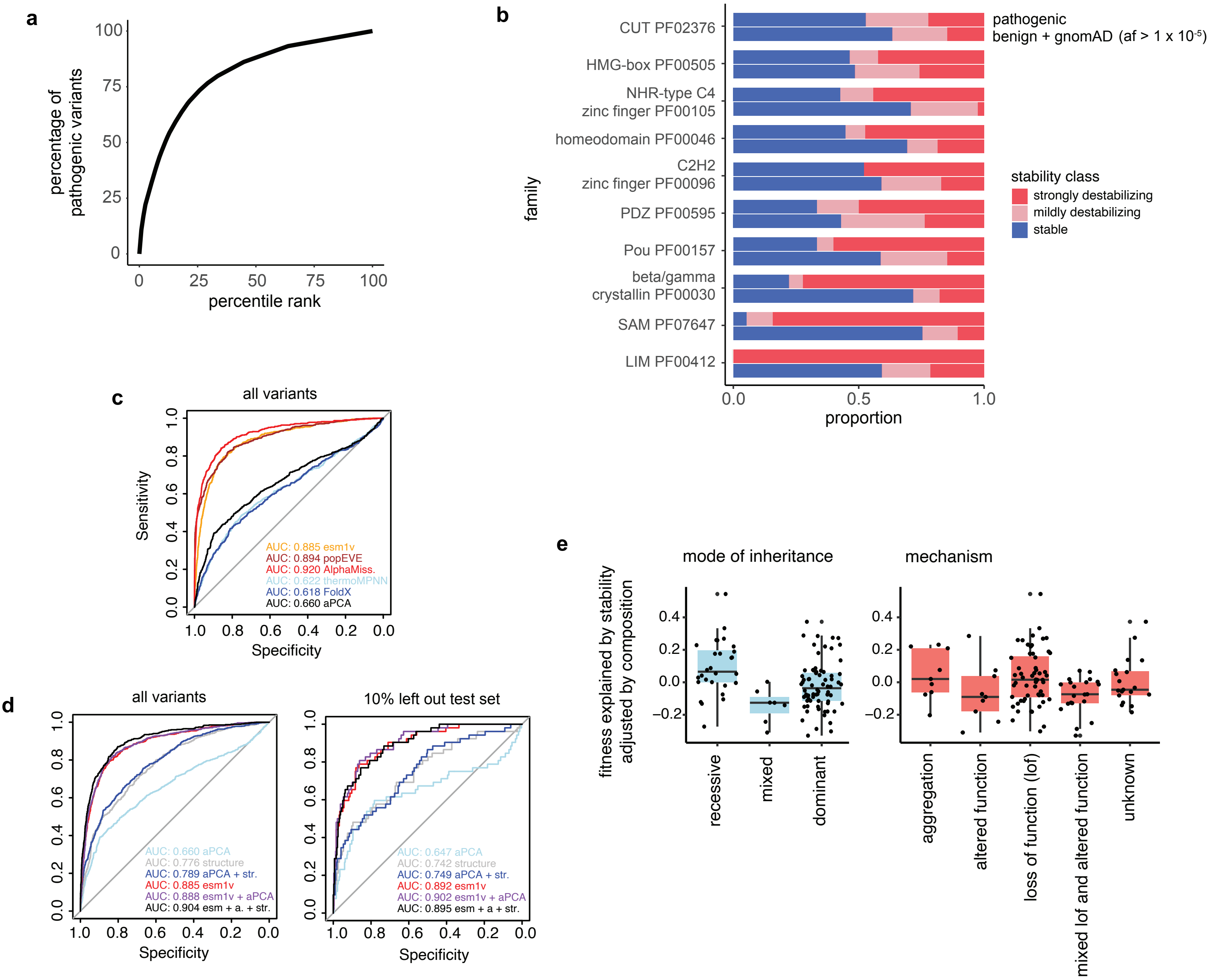


ED Figure 3

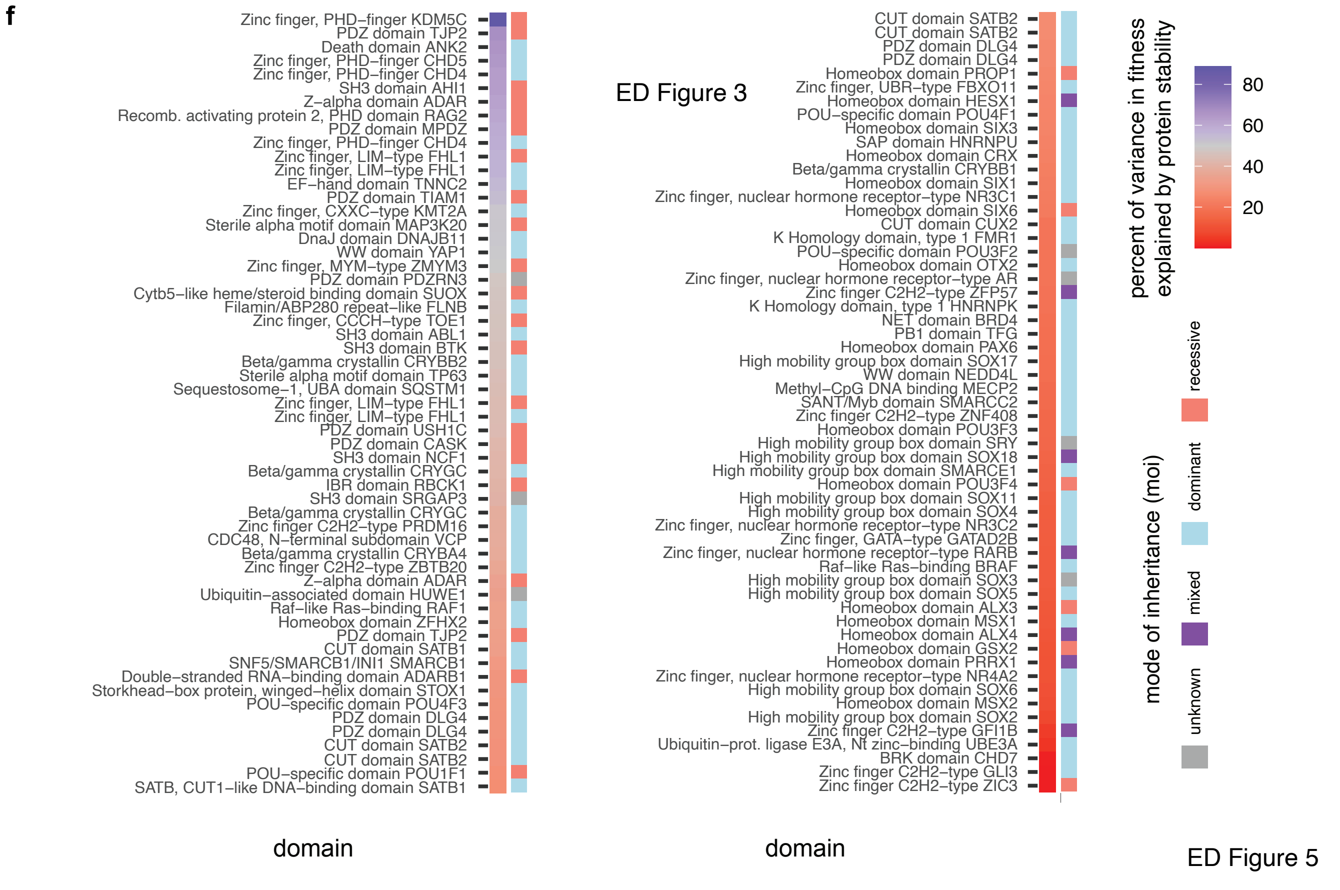


ED Figure 5

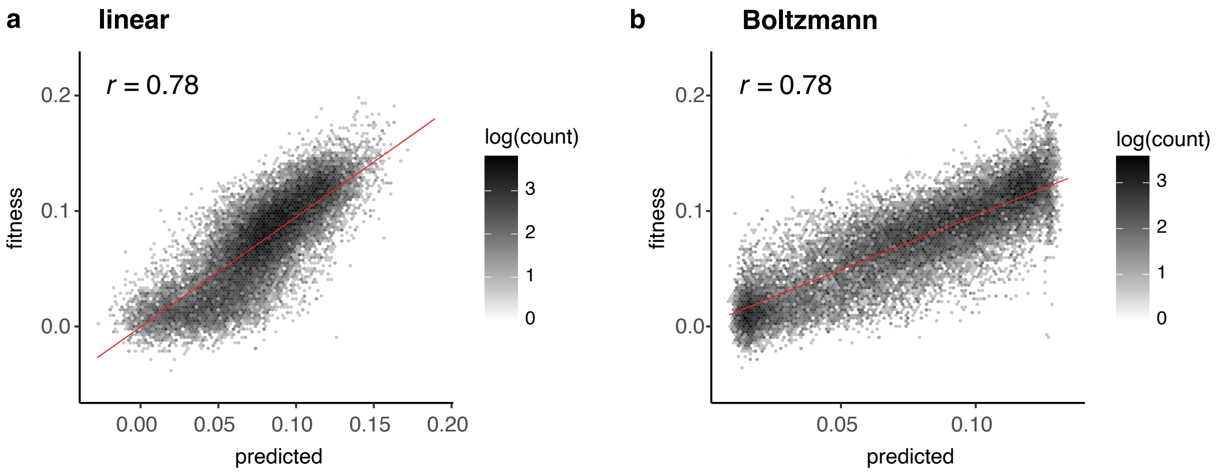
a
linear
r = 0.78
fitness
predicted
0.2
0.1
0.0
0.00
0.05
0.10
0.15
0.20
log(count)
3
2
1
0
b
Boltzmann
r = 0.78
fitness
predicted
0.2
0.1
0.0
0.05
0.10
log(count)
3
2
1
0

ED Figure 6

a

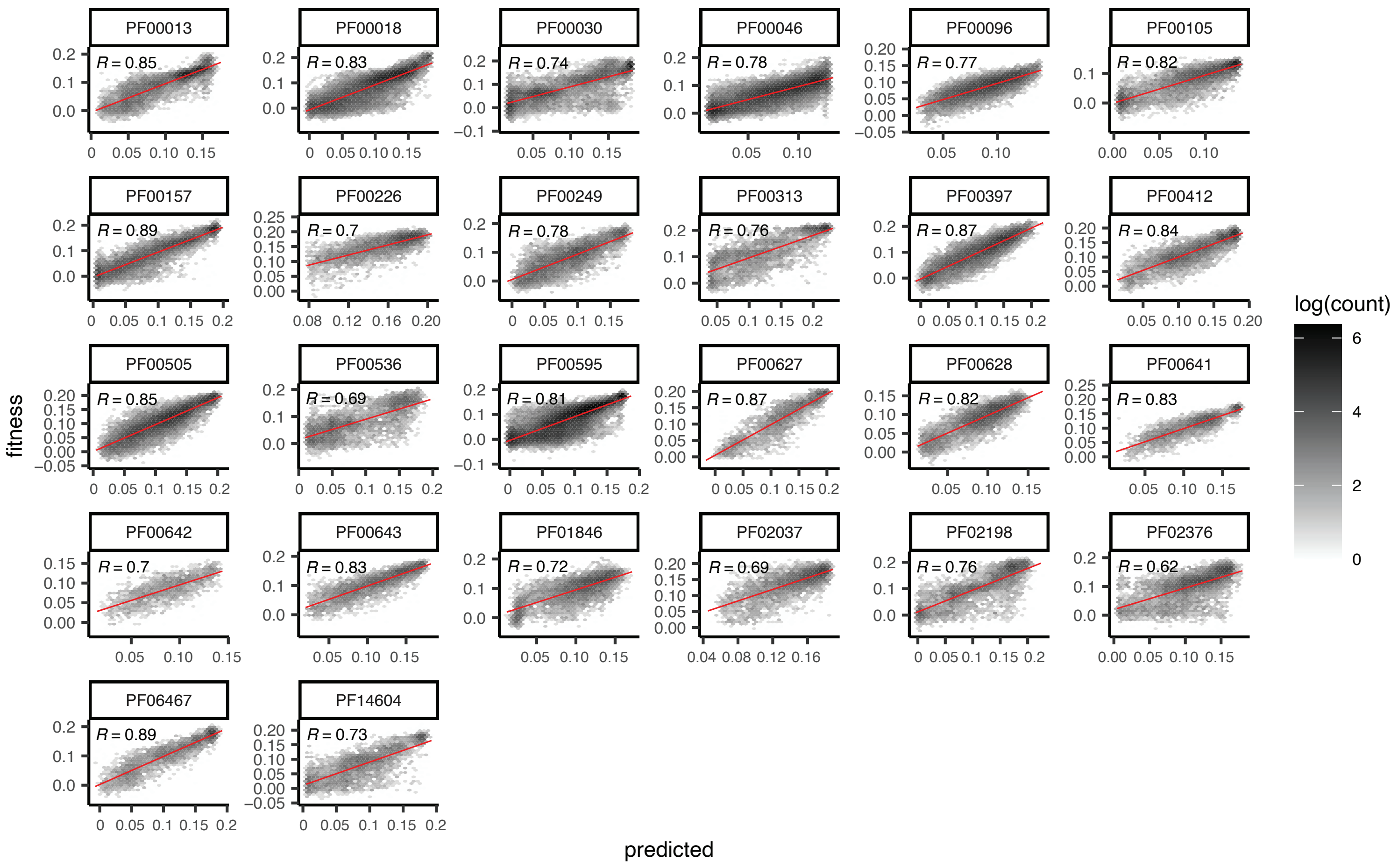


b

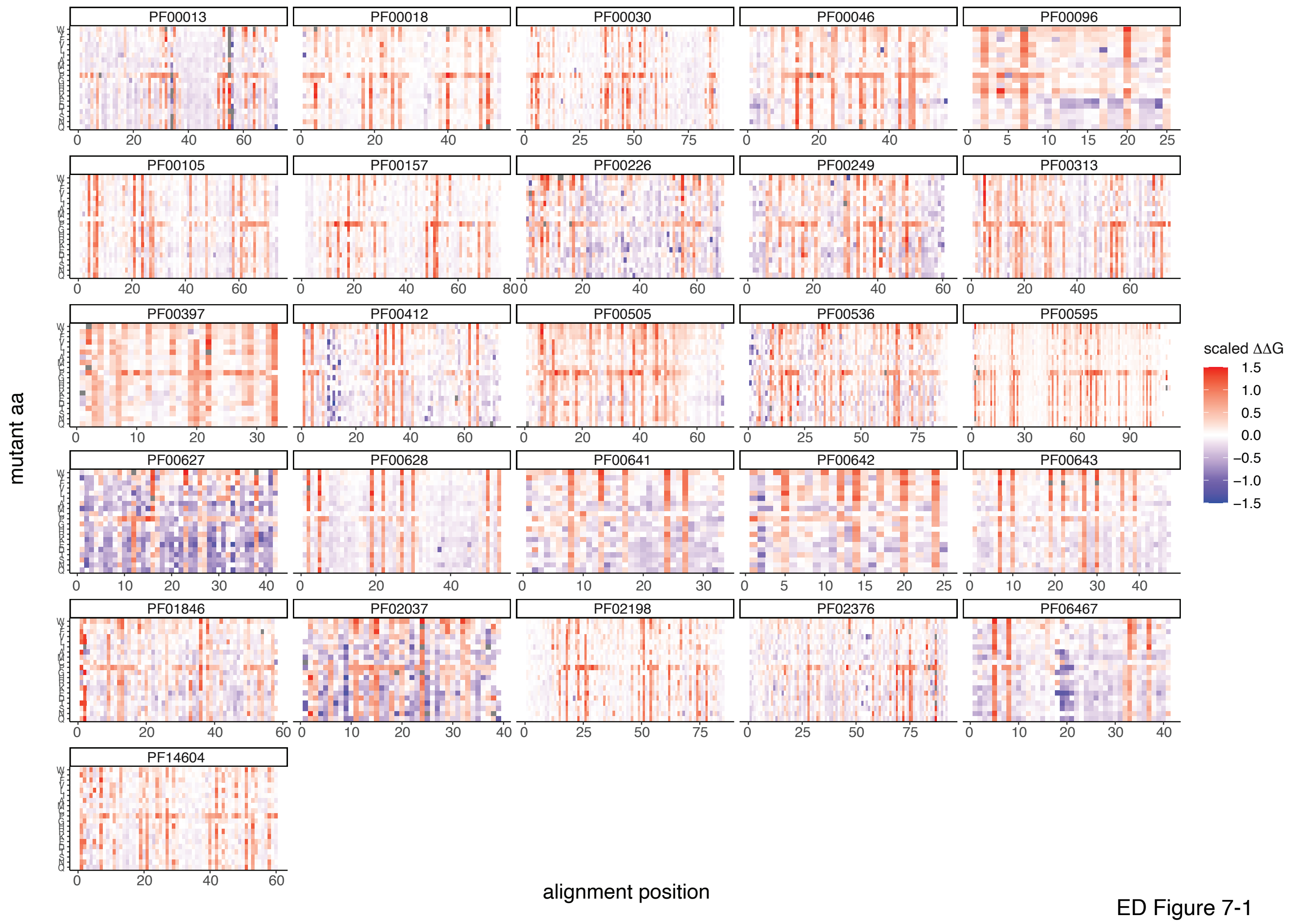


ED Figure 7-1

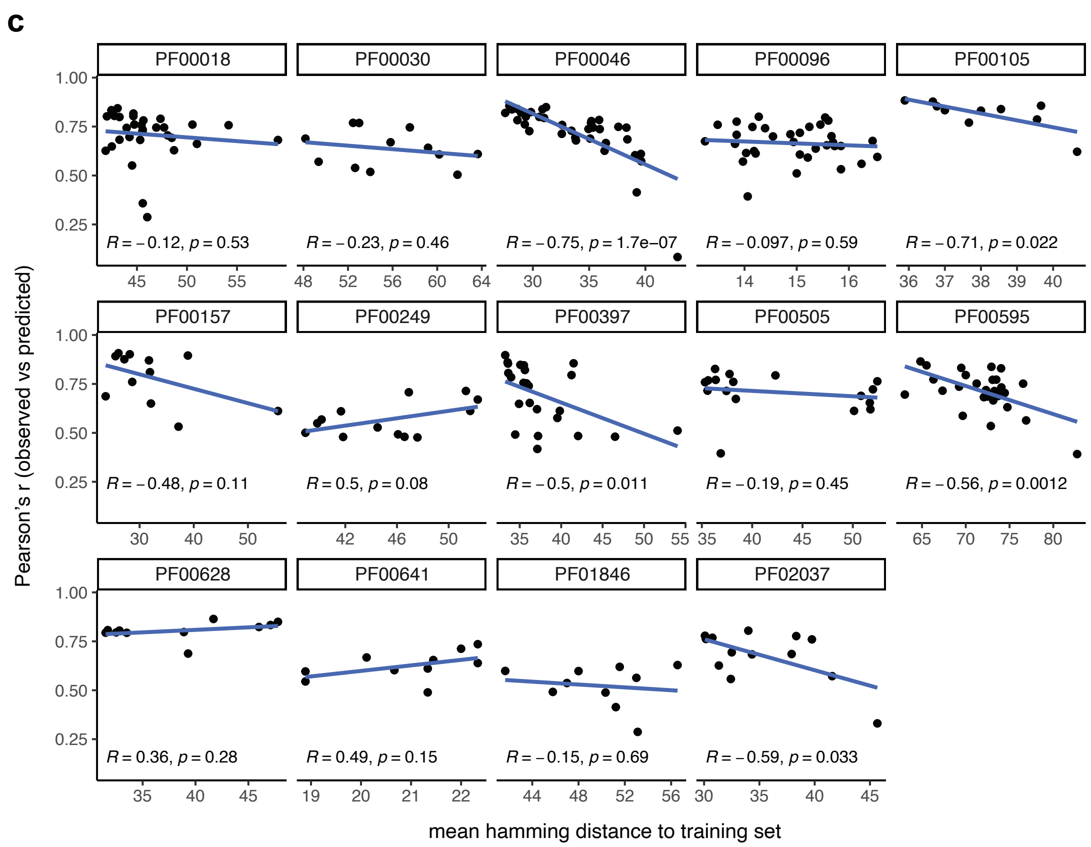

c
PF00018
PF00030
PF00046
PF00096
PF00105
PF00157
PF00249
PF00397
PF00505
PF00595
PF00628
PF00641
PF01846
PF02037
R = −0.12, p = 0.53
R = −0.23, p = 0.46
R = −0.75, p = 1.7e−07
R = −0.097, p = 0.59
R = −0.71, p = 0.022
R = −0.48, p = 0.11
R = 0.5, p = 0.08
R = −0.5, p = 0.011
R = −0.19, p = 0.45
R = −0.56, p = 0.0012
R = 0.36, p = 0.28
R = 0.49, p = 0.15
R = −0.15, p = 0.69
R = −0.59, p = 0.033
Pearson's r (observed vs predicted)
mean hamming distance to training set


ED Figure 7-2

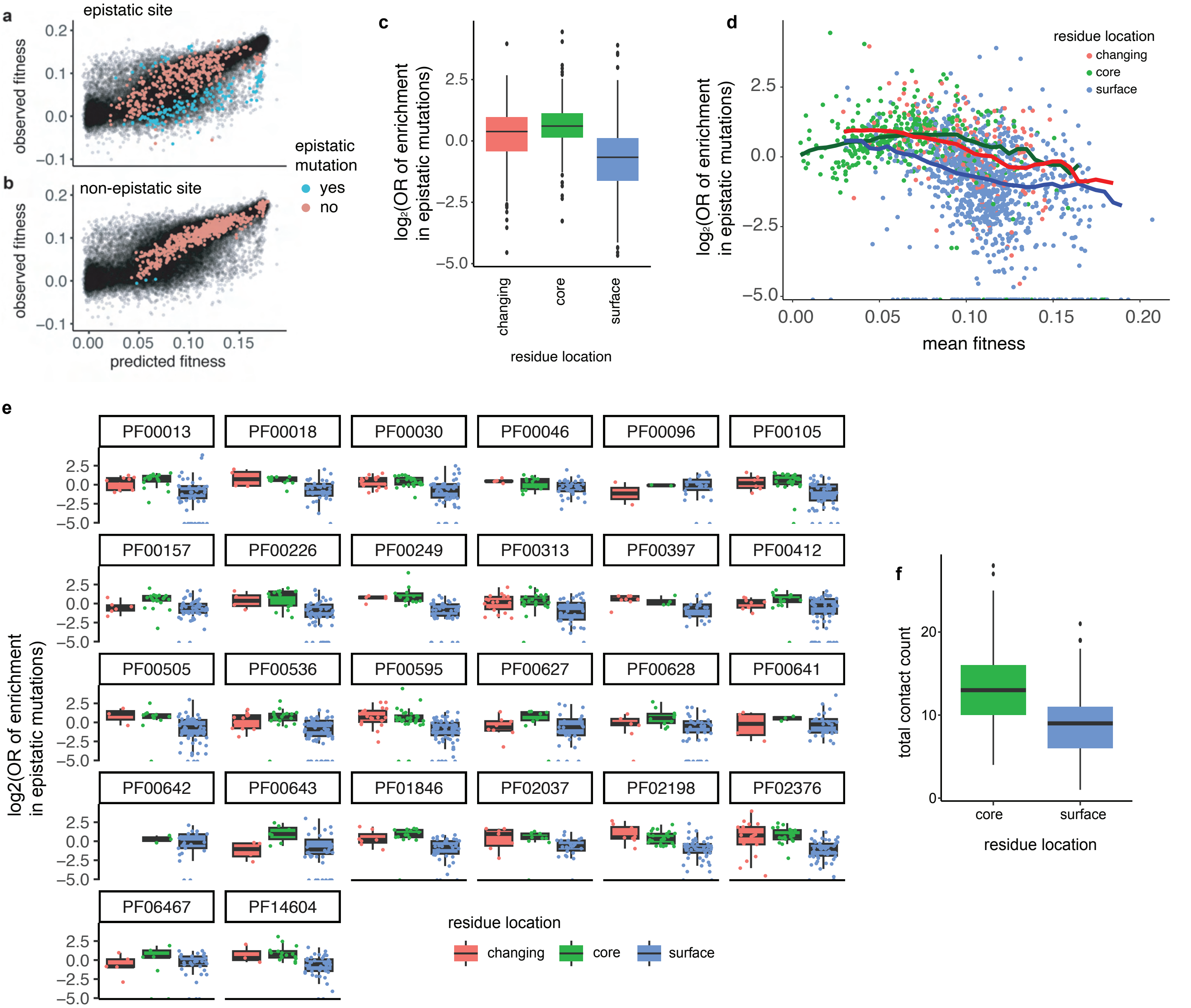

a
epistatic site
observed fitness
b
non-epistatic site
predicted fitness
epistatic mutation
yes
no
c
log₂(OR of enrichment in epistatic mutations)
changing
core
surface
residue location
d
residue location
changing
core
surface
mean fitness
e
PF00013
PF00018
PF00030
PF00046
PF00096
PF00105
PF00157
PF00226
PF00249
PF00313
PF00397
PF00412
PF00505
PF00536
PF00595
PF00627
PF00628
PF00641
PF00642
PF00643
PF01846
PF02037
PF02198
PF02376
PF06467
PF14604
log2(OR of enrichment in epistatic mutations)
residue location
changing
core
surface
f
total contact count
core
surface
residue location


ED Figure 8

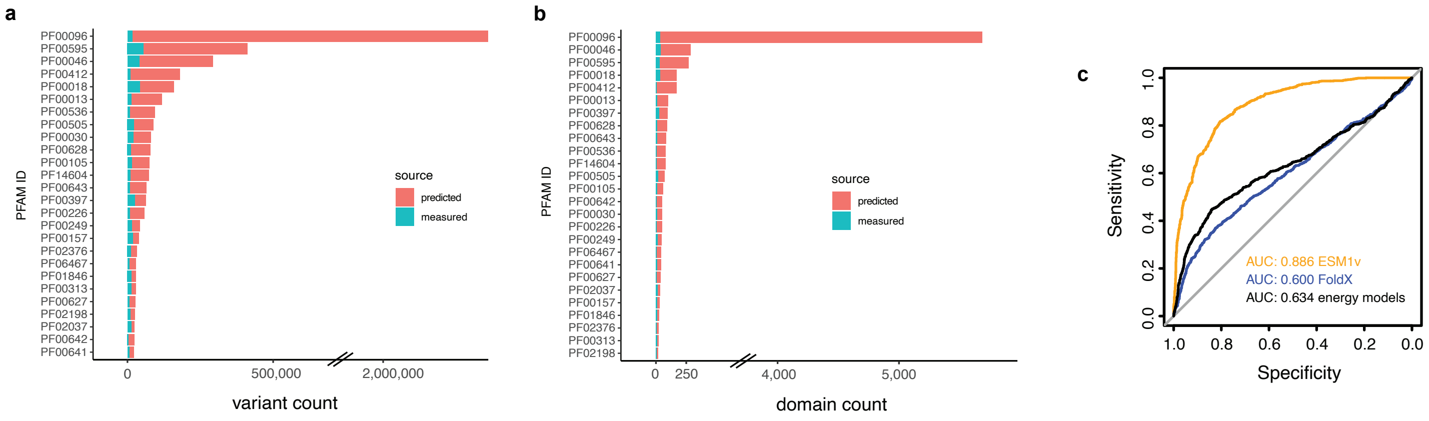
a
PFAM ID
PF00096
PF00595
PF00046
PF00412
PF00018
PF00013
PF00536
PF00505
PF00030
PF00628
PF00105
PF14604
PF00643
PF00397
PF00226
PF00249
PF00157
PF02376
PF06467
PF01846
PF00313
PF00627
PF02198
PF02037
PF00642
PF00641
source
predicted
measured
0
500,000
2,000,000
variant count
b
PFAM ID
PF00096
PF00046
PF00595
PF00018
PF00412
PF00013
PF00397
PF00628
PF00643
PF00536
PF14604
PF00505
PF00105
PF00642
PF00030
PF00226
PF00249
PF06467
PF00641
PF00627
PF02037
PF00157
PF01846
PF02376
PF00313
PF02198
source
predicted
measured
0
250
4,000
5,000
domain count
c
Sensitivity
Specificity
1.0
0.8
0.6
0.4
0.2
0.0
AUC: 0.886 ESM1v
AUC: 0.600 FoldX
AUC: 0.634 energy models

ED Figure 9
